# Evolutionary rates of nuclear and organellar genomes are linked in land plants

**DOI:** 10.1101/2024.08.05.606707

**Authors:** Yasmin Asar, Hervé Sauquet, Simon Y. W. Ho

**Author notes:** Corresponding author: Yasmin Asar **Email:**.

## Abstract

- Plants carry genetic material in three compartments, the nuclear, mitochondrial, and chloroplast genomes. These genomes interact with each other to various degrees and are subject to shared evolutionary drivers exerted by their host organisms. However, it is not clear whether the three plant genomes display covarying evolutionary signals.
- We tested for correlated evolutionary rates between nuclear and organellar genomes using extensive data sets from the major clades of land plants (Embryophyta), including mosses, ferns, gymnosperms, and angiosperms. To examine the evolutionary dynamics in parasitic angiosperms, which are under distinctive selective pressures, we analysed data sets from mistletoes, broomrapes, sandalwoods, and rafflesias.
- Evolutionary rates of nuclear and organellar genomes were positively linked in land plants, except in the parasitic angiosperms. We also found similar positive correlations for rates of nonsynonymous and synonymous change between nuclear and organellar genomes. Our results also reveal extensive evolutionary rate variation across land plant taxa.
- Overall, we find that nuclear, mitochondrial, and chloroplast genomes in land plants share similar drivers of mutaNon rates, despite considerable variaNon in life history, morphology, and genome sizes among clades. Our findings lay the foundaNon for further exploraNon of the impact of co-evoluNonary interacNons on shared evoluNonary rates between genomes.

## Introduction

Land plants (embryophytes) colonised the terrestrial environment between 583 and 459 Myr ago (Morris *et al*., 2018; Nie *et al*., 2020) and have diversified into an estimated 435,000 living species (Enquist *et al*., 2019), filling almost every ecological niche on Earth (Fig. 1). Whilst land plants display an immense range of morphological, ecological, and cellular differences, almost all have their genetic content carried in three separate genomes, found in their nuclei, mitochondria, and chloroplasts. Mitochondria are responsible for critical cell functions such as cellular respiration, immune function, and oxidative phosphorylation (Sloan *et al*., 2014; Zervas *et al*., 2019; Wang *et al*., 2022). Chloroplasts are primarily responsible for photosynthesis, transforming light and water into glucose (Martin & Herrmann, 1998). Unlike the nuclear genome, chloroplasts and mitochondrial genomes are much smaller, responsible for far fewer cell functions, effectively haploid, unable to sexually recombine, and are uniparentally transmitted, usually maternally (but see Greiner *et al.,* 2015 for exceptions).

**Fig. 1.**
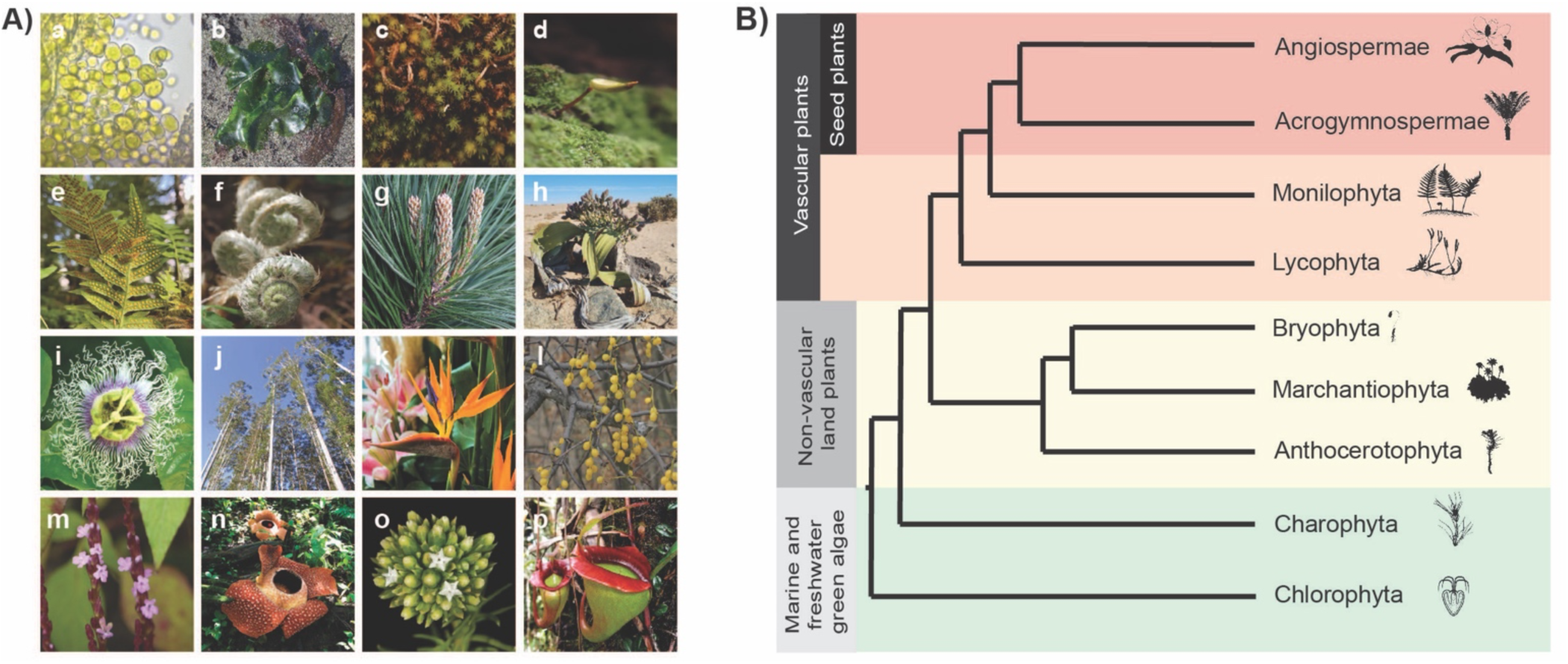
Morphological diversity and phylogeny of the green plants (Viridiplantae). (A) Representatives of the major clades of land plants that were analysed in this study (c–j) and close relatives of land plants, the chlorophyte green algae (a-b). a) *Chlorella* sp., b) *Ulva* sp., c) *Takakia* sp., d) *Buxbaumia aphylla*, e) *Polypodium glycyrrhiza*, f) *Polystichium acrostichoides*, g) *Pinus radiata*, h) *Welwitschia mirabilis*, i) *Passiflora edulis*, j) *Eucalyptus grandis*, k) *Strelitzia reginae*, l) *Loranthus europaeus*, m) *Striga gesneroides*, n) *Rafflesia arnoldii*, o) *Thesium scabrum*, and p) *Nepenthes jacquelinae*. All photographs were sourced from OpenVerse (https://openverse.org). Authors and Creative Commons licenses are supplied, unless image is in the public domain: a) Owen Strickland under CC BY 4.0, b) Ryan Hodnett under CC BY-SA 4.0, f) Derek Ramsey under CC BY-SA 2.5, g) Tony Wills under CC BY 2.5, h) Marc Fradera-Soler under CC BY 4.0, i) Sam Fraser-Smith under CC BY 2.0, j) Peter Woodard under CC BY-SA 3.0, k) fotopamas under CC BY 2.0, m) Dinesh Valke under CC BY-SA 2.0, n) SofianRafflesia under CC BY-SA 4.0, o) SAplants under CC BY-SA 4.0, and p) Shawn Mayes under CC BY-SA 3.0. (B) Simplified phylogeny of Viridiplantae (the green plants). Relationships are summarised from those described in One Thousand Plant Transcriptomes Initiative (2019). Plant silhouettes are in the public domain and available at http://www.phylopic.org.

Considerable evolutionary and structural variation can be seen across the nuclear genomes of land plants, with extensive degrees of genome duplication and polyploidy. For instance, the largest known genome (1*C* = 160.45 Gb) occurs in an octoploid New Caledonian fork fern, *Tmesipteris oblanceolata* (Fernández *et al*., 2024). Polyploidy is likely to have provided an important foundation for the phenotypic novelties that allowed the flowering plants to radiate and speciate (Soltis *et al*., 2019; Benton *et al*., 2022). Plant mitochondrial genomes share the same endosymbiotic origin, cellular function, and uniparental transmission as those of animals, yet they exhibit a range of structures, from circular to branched, are much larger (0.2–20 Gb), and typically have very low rates of nucleotide substitution (Wolfe *et al*., 1987; Drouin *et al*., 2008; Chen *et al*., 2017; Morley & Nielsen, 2017). In contrast, chloroplast genomes are remarkably conserved in size, structure, and content across land plants. They have much lower substitution rates than nuclear genomes in land plants (Wolfe *et al*., 1987; Drouin *et al*., 2008; Smith, 2015). Chloroplast genomes are circular, 110– 200 kb in length, and include 80–90 protein-coding genes, four ribosomal RNA genes, and 30–31 transfer RNA genes (Kaur *et al*., 2022). However, there are massive reductions in plastome content in parasitic angiosperms (Graham *et al*., 2017; Zervas *et al*., 2019). This reduction reaches its most extreme in the family Rafflesiaceae, where two species have lost their entire chloroplast genome (Molina *et al*., 2014; Cai *et al*., 2021). Parasitic flowering plants are ‘outliers’ to the remaining land plants in other aspects; for instance, there are vastly accelerated rates of mitochondrial evolution along with gene loss in *Viscum* (mistletoes) (Havird *et al*., 2019; Zervas *et al*., 2019).

Although there are great differences in size, structure, and content between nuclear and organellar genomes, there are well-documented interactions between these genomic compartments. Coordinated change between the nucleus and mitochondria may be integral for survival; this manifests as mitonuclear covariation in animals, where the accumulation of deleterious mutations in mitochondrial genomes is compensated for by mutations in the nucleus (Martin & Herrmann, 1998; Weaver *et al*., 2022). The mitochondria have protein complexes that consist of subunits variously encoded by the mitochondrial and nuclear proteins, and these subunits need to interact properly to ensure full function (Martin & Herrmann, 1998; Weaver *et al*., 2022). This is particularly evident in males, where mutations that impede fertility in males are non-functional in females, and are retained in the population despite their deleterious effects (David *et al*., 2022). Furthermore, hybrids in various animals show reduced fitness because of incompatibilities between maternally inherited mitochondrial genes and paternally inherited nuclear-encoded mitochondrial proteins (Burton *et al*., 2013; Sloan *et al*., 2017). Thus, mitonuclear covariation can act as a post-zygotic barrier on the path to speciation (Greiner *et al*., 2015; Hill, 2016). This is also demonstrated in hybrid plants, where expression of chimeric open reading frames in the mitochondria causes cytoplasmic male sterility, which results in a failure to produce functional pollen (Bentolila *et al*., 2002; Delph *et al*., 2007; Fujii *et al*., 2011). Only when there are compensatory changes in nuclear-encoded proteins can there be restoration of male fertility.

Given the co-evolutionary interactions between the nuclear and organellar genomes, we would expect their evolutionary rates to show some level of association. Such rate correlations have been observed in the nuclear and mitochondrial genomes of crustaceans, bivalves, mammals, yeast, and insects (Barreto *et al*., 2018; Yan *et al*., 2019; Piccinini *et al*., 2021; Formaggioni *et al*., 2022; Weaver *et al*., 2022; Biot-Pelletier *et al*., 2023; Tao *et al*., 2024). However, there has been mixed evidence of evolutionary rate correlations across nuclear and organellar genomes in plants, with some researchers discounting the existence of coevolutionary interactions between nuclear and chloroplast genomes (Hill, 2020). Recent studies have found strong signals for rate correlations in data sets comprising hundreds of nuclear and chloroplast genes from 20–25 angiosperm taxa (Williams *et al*., 2019; Forsythe *et al*., 2021), as well as evidence for linkage disequilibrium in nuclear and mitochondrial genes across 3439 genomes of seven angiosperm species (Lian *et al*., 2024). Similar correlations have been detected in studies of individual genera and families of angiosperms (Zhang *et al*., 2015; Rockenbach *et al*., 2016; Yang *et al*., 2023; Khachaturyan *et al*., 2024). Neverthless, the amount of co-ordination between nuclear, mitochondrial, and chloroplast genomes has not been fully explored across the diversity of land plants. This leaves uncertainty about whether the interactions between the three genomes have left broad-scale signatures of covariation in their evolutionary rates.

In this study, we characterise evolutionary rate variation and test for correlations in rates of nuclear, mitochondrial, and chloroplast genome evolution using extensive data sets from major lineages of land plants. We first conduct a simulation study to validate the performance of three phylogenetic methods used to test for evolutionary rate correlations. We then use these methods to analyse data sets of mosses, ferns, gymnosperms, and angiosperms. Our analyses reveal strong evolutionary rate correlations between nuclear, mitochondrial, and chloroplast genes across major lineages of land plants. In contrast, these patterns are not detected in parasitic angiosperms, perhaps a consequence of their unique habit and reduced genomes.

## Materials and Methods

### Data collection and curation

We collected nucleotide sequence data for nuclear, mitochondrial, and chloroplast markers from a diverse range of plant taxa, including mosses, ferns, gymnosperms, angiosperms, as well as a number of genera and families of parasitic angiosperms. We use the formal terminology, as per Cantino *et al*. (2007) and Bechteler *et al*. (2023). Thus, ‘mosses’ refers to Bryophyta, ‘ferns’ refers to Monilophyta, ‘gymnosperms’ refers to the extant Acrogymnospermae, and ‘angiosperms’ to Angiospermae. Full details of the data sets and their sources are provided in Supporting Information (Table S1).

We targeted the most recent genomic studies, which included all described protein-coding genes for the organellar data sets, with taxon sampling to the greatest extent achievable under our computational limits. Overall, for the four major land plant clades (mosses, ferns, gymnosperms, and angiosperms), we assembled data with mean alignment lengths of 167,688 bp for nuclear DNA, 27,341 bp for mitochondrial DNA, and 60,574 bp for chloroplast DNA. The average number of taxa that were used for these comparisons was *n* = 86. Gene alignments were smaller for data sets of parasitic angiosperms, with mean lengths of 3,298 bp, 5,585 bp, and 51,230 bp, for nuclear, mitochondrial, and chloroplast data sets respectively (see Supporting Information Table S1 and Notes S1). The average number of taxa for these comparisons was *n* = 70.

We pruned the nuclear and organellar data sets to ensure that taxa matched between the data sets. We then inferred phylograms using maximum likelihood in IQ-TREE2 (version 2.2.2.6; Bui et al. 2020) and Bayesian inference in BEAST2 (version 2.6.6; Bouckaert et al. 2019). For all analyses, tree topologies were fixed to match those from the source studies (Supporting Information Table S1). We did not allow for the possibility that separate genomic compartments can support highly divergent phylogenies, but topological discordance is a source of noise that would weaken any signals of evolutionary rate correlation.

### Phylogenetic analyses

We inferred phylograms using maximum-likelihood analysis. When alignments of individual genes were available, we selected the partitioning scheme and nucleotide substitution models in IQ-TREE2 (-m TESTMERGEONLY). For gene data sets with sequences that were well aligned and in frame, we also split the sequences into 1st+2nd and 3rd codon sites. We used these as proxies for nonsynonymous and synonymous sites, respectively, allowing us to test for correlations at these sites separately. Branch lengths were proportionally linked between partitions. We constrained the tree topology (-g) and inferred branch lengths for each data set.

We then inferred trees using Bayesian analysis in BEAST2, using the same partitioning schemes as in the maximum-likelihood analyses. For each partition, the substitution model was averaged using bModeltest (Bouckaert & Drummond, 2017). Uncorrelated lognormal relaxed clock models were unlinked between genomes (nuclear, chloroplast, and mitochondrial), with one clock model having a mean rate of 1.0 and with the other clock model having a mean rate that was inferred in the analysis (Drummond *et al*., 2006). The mean rate of the clock model was assigned a uniform prior with bounds of 0 and 1,000,000. To allow rate variation across partitions, each was assigned its own relative rate parameter. The tree topology was fixed, as in the maximum-likelihood analyses, with a birth-death tree prior on the relative node times.

Each analysis was run for at least 100,000,000 MCMC steps, with sampling every 10,000 steps. The sampling frequency was varied according to the chain length, with at least 10,000 trees sampled from the posterior for each run. At least three independent runs were completed for each analysis and their samples combined in LogCombiner, with at least 10% of each run discarded as burn-in. Effective sample sizes, checked in Tracer (Rambaut *et al*., 2018) were greater than 200 for almost all parameters, including the likelihood and posterior. The combined tree samples were summarised using TreeAnnotator.

### Methods for testing evolutionary rate correlations

We validated the methods for detecting correlated evolutionary rates by ensuring that they were both accurate (with low rate of false-positive detection) and powerful (able to detect linked rates of evolution across a variety of settings). We performed a comprehensive simulation study, described in the Supporting Information (see Methods S1 and Fig. S1–S6), to evaluate methods based on correlations of (*i*) root-to-tip distances, (*ii*) independent sister-pair contrasts, and (*iii*) branch rates inferred using Bayesian relaxed-clocks. All three methods were found to be accurate, powerful, and fit for use when testing for positive evolutionary rate correlations. These results are consistent with those from our previous evaluation of these methods in detecting evolutionary rate correlations between molecular and morphological data sets (Asar *et al*., 2023).

Root-to-tip distances were calculated using the package *adephylo* in R. The outgroup taxa were removed from the nuclear, mitochondrial, and chloroplast phylograms inferred in our maximum-likelihood analyses above, then patristic distances between the root of the tree to each tip were calculated. We then tested for pairwise correlations between the root-to-tip distances of the phylograms. We used a permutation correlation test (perm.cor.test function in *jmuOutlier*) (Higgins, 2004) using 200,000 simulations (one-sided), implemented in R (Garren, 2017), to avoid violations caused by use of non-independent data points.

The second method of testing for correlated rates involved taking phylogenetically independent pairs of taxa (sister-pairs) and comparing their relative branch lengths between nuclear, mitochondrial, and chloroplast phylograms. Using the R package *diverge* (Anderson & Weir, 2020), we extracted sister species that shared a most recent common ancestor to the exclusion of other sister pairs, which avoids the problem of phylogenetic non-independence but reduces the amount of data (Felsenstein, 1985; Bromham *et al*., 2002). We computed the difference between the branch lengths of sister taxa for each of the plant organellar phylograms. We tested for positive correlations between these contrasts using the non-parametric Spearman’s rank correlation test, to allow for violations of bivariate normality and homoscedasticity. We log-transformed and standardised all contrasts by dividing them by the square root of the sum of branch lengths (Garland *et al*., 1992; Freckleton, 2000; Welch & Waxman, 2008; Lanfear *et al*., 2010). After investigating whether the data met the assumptions of the independent sister-pairs contrasts method, we used a two-sample sign test where the assumptions were violated. See Supporting Information for full description of assumption tests and their results for all comparisons (Methods S2, Table S8–S16).

The third method of testing for correlated rates was based on the inferred posterior mean branch rates from Bayesian phylogenetic analyses. We tested for positive correlations between the branch rates inferred using the uncorrelated lognormal relaxed clock for the plant nuclear, mitochondrial, and chloroplast data sets. We obtained the posterior mean branch rates in R, using the package *treeio* (version 1.22.0) (Wang *et al*., 2020). To compute the significance of the correlation, we used Spearman’s rank-order correlation test.

## Results

### Evolutionary rates across land plants

Our analyses revealed considerable evolutionary rate variation in nuclear and chloroplast genomes across the major lineages of land plants (Fig. 2A): mosses, ferns, gymnosperms, and angiosperms. We estimated evolutionary rates by dividing the mean root-to-tip distance of each phylogenetic tree by the estimated age of its root node (Supporting Information Table S2). Given that the age of the angiosperms continues to be debated, with molecular estimates ranging from the Late Jurassic to the Permian (Sauquet *et al*., 2022), we used multiple estimates for the crown-age of this group (Fig. 2A, Supporting Information Table S2). Estimated evolutionary rates of nuclear genomes ranged from 4.46×10^-4^ substitutions/site/Myr in acrogymnosperms to 2.16×10^-3^ substitutions/site/Myr in angiosperms. In contrast, the estimated evolutionary rate of chloroplast genomes varied from 4.56×10^-4^ substitutions/site/Myr in acrogymnosperms to 9.67×10^-4^ substitutions/site/Myr in ferns. Rates of mitochondrial genome evolution were substantially lower than those of nuclear and chloroplast genomes, ranging from 2.33×10^-4^ substitutions/site/Myr in mosses to 4.69×10^-4^ substitutions/site/Myr in angiosperms. All data sets of land plant nuclear, mitochondrial, and chloroplast genomes used in analyses are described in Supporting Information Table S1.

**Fig. 2.**
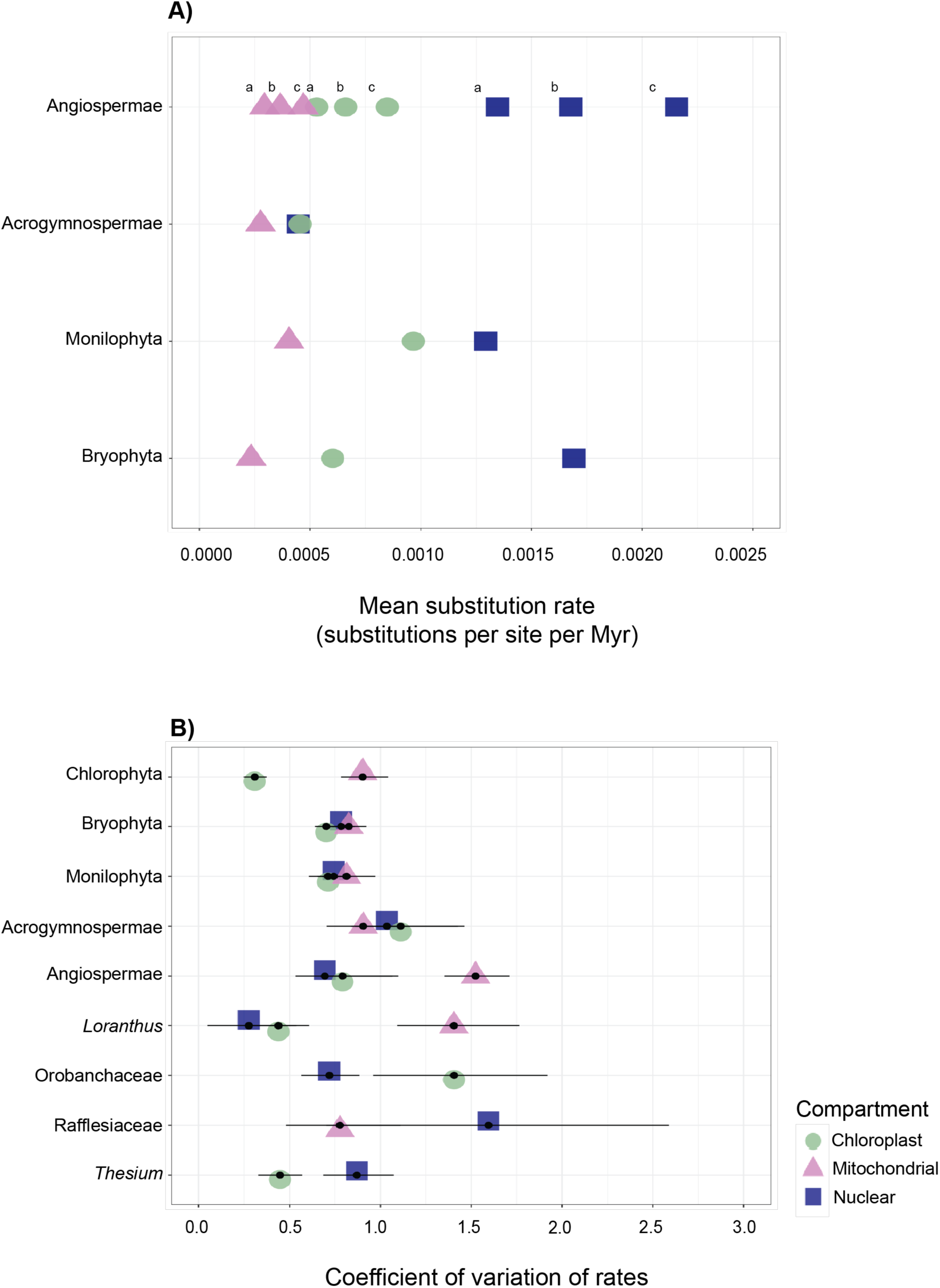
There is pronounced diversity in evolutionary rates and among-lineage rate heterogeneity amongst the major lineages of land plants. (A) Estimates of evolutionary rates for nuclear markers (dark blue squares), mitochondrial markers (violet triangles), and chloroplast markers (green circles) of the major embryophyte lineages. Three estimates of evolutionary rates are provided for Angiospermae, depending on whether a Middle Triassic ∼246 Ma (a), Early Jurassic ∼198 Ma (b), or Late Jurassic ∼ 154 Ma (c) age of origin is assumed. These age estimates are from the analyses using constrained, relaxed, and unconstrained complete calibrations reported by Ramírez-Barahona *et al.,* (2020). Plant silhouettes are in the public domain and available at http://www.phylopic.org. (B) Coefficient of variation of rates, a measure of among-lineage rate heterogeneity, estimated for each of the plant clades in this study. The nuclear, mitochondrial, and chloroplast genomic compartments are represented by dark blue squares, violet triangles, and green circles, respectively. Estimates were obtained by Bayesian relaxed-clock analysis and are shown as posterior means accompanied by 95% credibility intervals.

Across four major lineages of land plants, we identified disparate levels of among-lineage rate variation as measured by the coefficient of variation of rates (σ) (Fig. 2B). This metric was calculated in a Bayesian phylogenetic analysis by dividing the standard deviation of the branch rates by the mean of the branch rates, with values close to zero indicating clocklike evolution (Drummond *et al*., 2006). In the major lineages of land plants, the nuclear and chloroplast data sets had similar levels of moderate rate heterogeneity, with mean σ ranging from 0.694 to 1.11 across all groups (Fig. 2B; Supporting Information Table S3). In angiosperms, the mitochondrial mean σ was 1.52 (95% CI 1.35– 1.71), reflecting an extreme degree of rate variation among lineages. Among parasitic angiosperms, σ was especially pronounced in nuclear genes of Rafflesiaceae (σ = 1.60, 95% CI 0.702–2.59), mitochondrial genes in *Loranthus* (σ = 1.40, 95% CI 1.09–1.76), and chloroplast genes in Orobanchaceae (σ = 1.41, 95% CI 0.961–1.92). Concordant results were obtained when we estimated the degree of among-lineage rate heterogeneity using the coefficient of variation of root-to-tip distances in maximum-likelihood phylogenetic trees (Supporting Information Fig. S7, Table S3).

### Evolutionary rate correlations between genomes

We tested for evolutionary rate correlations across genomic compartments of the major land plant lineages using available genomic data sets (Supporting Information Table S1). We began by conducting a comprehensive simulation study to validate three methods of analysis (described fully in Supporting Information Methods S1): (*i*) correlations of root-to-tip distances, based on maximum-likelihood phylograms; (*ii*) independent sister-pair contrasts, based on maximum-likelihood phylograms; and (*iii*) correlations of branch rates inferred using Bayesian relaxed-clocks. We found that all three methods were accurate, with false positives detected across an average of 3.8% of replicates, and that the methods were moderately powerful, with the average detection of true correlations at 88.5% (Supporting Information Fig. S1–S6). The independent sister-pair contrasts method was the most conservative, being unable to detect any rate correlations when there were only 18 ingroup taxa in the simulations. In these cases, there were only a small number of sister-pair contrasts (*n*=3) (Supporting Information Fig. S1, S5). When the 18-taxon simulation setting was excluded for independent sister-pair contrasts, the average detection of true correlations overall increased to 99.5% across the three methods. Accordingly, we focused on sampling data sets that had more than 20 ingroup taxa (Supporting Information Table S1). We found that the third method, correlations of branch rates inferred using Bayesian relaxed-clocks, had the greatest accuracy and power (Supporting Information Fig. S2, S6). Thus, we focus on the results obtained using this method, although the two other methods yielded concordant results (Supporting Information Fig. S8–S10).

We found consistently strong, significant correlations between evolutionary rates of nuclear and chloroplast genomes (Fig. 3; *r_s_* ≥ 0.48 and *p* ≤ 10^-8^ for all comparisons). This result was found across all of the major land plant lineages that we assessed. Similarly, there was a significant association between evolutionary rates of nuclear and mitochondrial genomes (Supporting Information Fig. S8; *r_s_* ≥ 0.25 and *p* ≤ 10^-3^) and between almost all mitochondrial and chloroplast genomes (Supporting Information Fig. S8; *r_s_* ≥ 0.28 and *p* < 0.01). All tested associations were significant, with the exception of angiosperm mitochondrial and chloroplast genomes, which was weakly positive but statistically insignificant (*r_s_* = 0.032, *p* = 0.27). We were also unable to detect an evolutionary rate correlations between mitochondrial and chloroplast genomes in a clade of marine algae, closely related to land plants, Chlorophyta (Supporting Information Fig. S11, *r_s_* = 0.18 and *p* = 0.13).

**Fig. 3.**
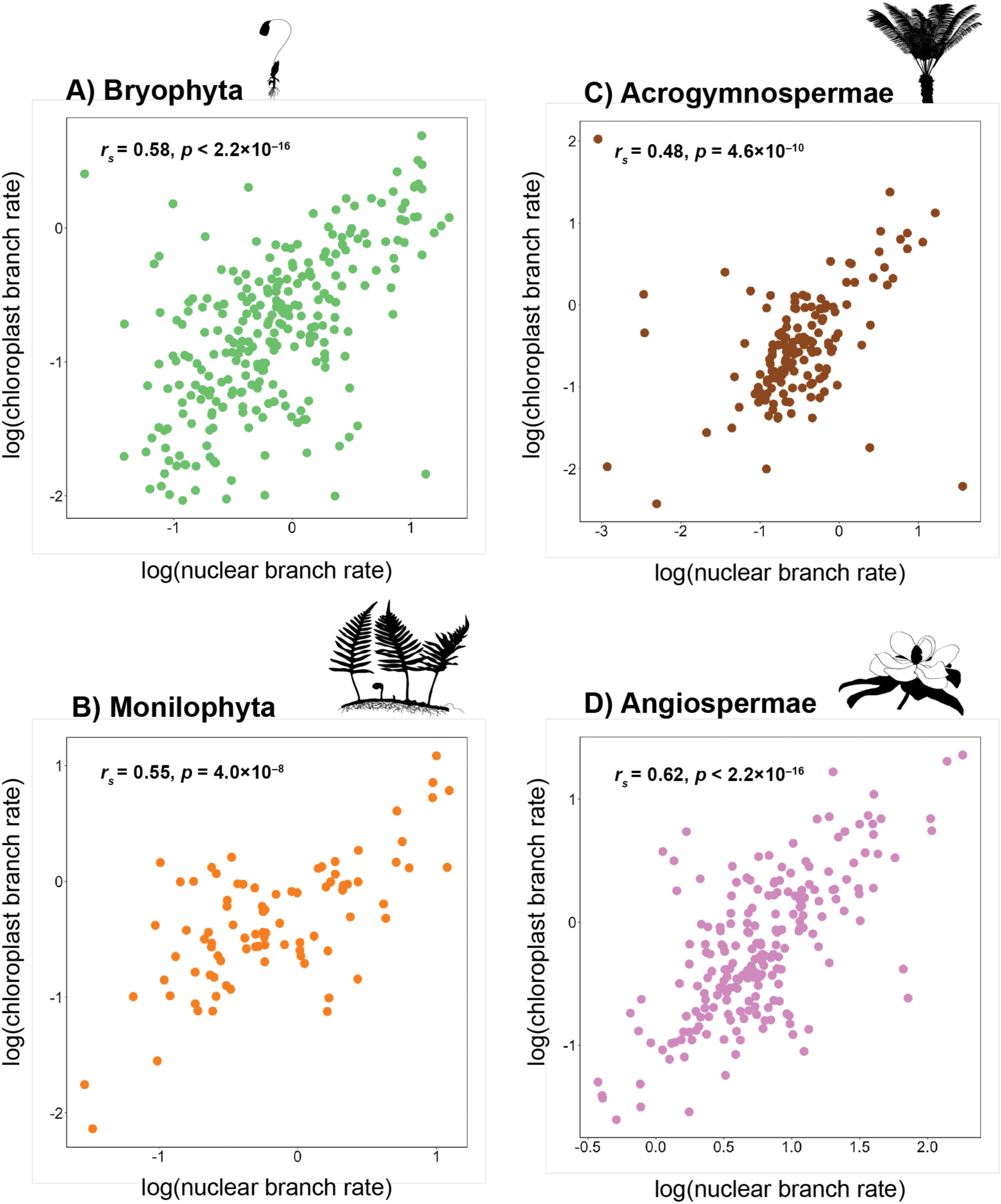
Rates of nuclear and chloroplast evolution are strongly linked in the major embryophyte clades. Log-transformed relative branch rates plotted for nuclear genome against the chloroplast genome for the major land plant clades. Branch rates were estimated using Bayesian relaxed-clock analysis. The correlation coefficient (*r_s_*) and *p*-value (calculated using Spearman’s rank correlation test) are shown for each plot. Bold font indicates *p*-values below 0.05. Plant silhouettes are in the public domain and available at http://www.phylopic.org.

To gain additional insight into the nature of the rate correlations, we performed further analyses of protein-coding sequences by separating the 1st+2nd and 3rd codon sites. We treated these as proxies for nonsynonymous and synonymous evolutionary changes, respectively. These analyses were possible for three data sets: mosses, ferns, and angiosperms. For both nonsynonymous and synonymous rates, we detected correlations between nuclear and mitochondrial genomes (Table 1; *r_s_* ≥ 0.12 and *p* ≤ 0.024) and between nuclear and chloroplast genomes (Table 1; *r_s_* ≥ 0.51 and *p* ≤ 0.01). Similarly, almost all comparisons of evolutionary rates between mitochondrial and chloroplast genomes detected significant associations (Table 1). However, we were unable to detect a correlation in nonsynonymous rates (*r_s_* = 0.011 and *p* = 0.42) and synonymous rates (*r_s_* = 0.033 and *p* = 0.27) between the mitochondrial and chloroplast genomes of angiosperms, despite the large sample size (*n* = 185). The results were concordant with those obtained using the two other methods that we employed (Supporting Information Table S4–S5). The ability to detect significant associations was reduced when we used sister-pair contrasts, as expected from the results of our simulations. However, all tested comparisons were significant when using root-to-tip distances, which was not unexpected, given the slightly inflated detection of rate correlations when using this method shown in our simulations (Supporting Information Methods S1, Fig. S2).

**Table 1.** Correlations between rates at 1^st^ + 2^nd^ sites and 3^rd^ sites for each of the organellar compartments in the major embryophyte clades. For each organellar comparison, the correlation coefficient (*r_s_*) and p-values (*p*) for a Spearman rank correlation test are recorded. Bold font indicates *p*-values below 0.05.

| Taxon<sup>a</sup> |  | Nuclear–Chloroplast |  | Nuclear–Mitochondrial |  | Chloroplast–Mitochondrial |  |
| --- | --- | --- | --- | --- | --- | --- | --- |
|  |  | 1st + 2nd | 3rd | 1st + 2nd | 3rd | 1st + 2nd | 3rd |
| Bryophyta | $r_s$ | 0.51 | 0.61 | 0.12 | 0.23 | 0.31 | 0.26 |
| | $p$ | **<2.2×10<sup>-16</sup>** | **<2.2×10<sup>-16</sup>** | **2.4×10<sup>-2</sup>** | **9.4×10<sup>-5</sup>** | **8.3×10<sup>-8</sup>** | **7.4×10<sup>-6</sup>** |
| Monilophyta | $r_s$ | 0.57 | 0.52 | 0.29 | 0.39 | 0.27 | 0.40 |
| | $p$ | **9.9×10<sup>-9</sup>** | **2.3 ×10<sup>-7</sup>** | **3.2×10<sup>-3</sup>** | **1.2×10<sup>-4</sup>** | **3.6×10<sup>-3</sup>** | **2.2×10<sup>-5</sup>** |
| Angiospermae<sup>b</sup> | $r_s$ | 0.40 | 0.30 | - | - | 0.011 | 0.033 |
| | $p$ | **1.1×10<sup>-3</sup>** | **1.0×10<sup>-2</sup>** | - | - | 4.2×10<sup>-1</sup> | 2.7×10<sup>-1</sup> |

<sup>a</sup>Number of taxa ( $n$ ) included for the comparisons was: $n = 135$ (Bryophyta), $n = 44$ (Monilophyta), $n = 51$ (Monilophyta, chloroplast–mitochondrial), $n = 30$ (Angiospermae, nuclear–chloroplast), and $n = 185$ (Angiospermae, chloroplast–mitochondrial).
<sup>b</sup>These comparisons refer to the data sets sourced from Zeng et al., (2014), Ruhfel et al., (2014), and Tyszka et al., (2023). This differs from the nuclear and chloroplast data sets presented in the main figure since these did not have site partitioning data available.

### Uncoupled evolutionary rates in parasitic flowering plants

We tested for evolutionary rate correlations across the nuclear, mitochondrial, and chloroplast genomes of four groups of parasitic angiosperms. For these analyses, we targeted the most recent genomic or multilocus studies with large taxon sampling (full description in Supporting Information Table S1 and see Notes S1). Our analyses failed to detect any associations in evolutionary rates between nuclear and chloroplast genomes in Orobanchaceae, *Thesium*, and *Loranthus*, and between nuclear and mitochondrial genomes in Rafflesiaceae (Fig. 4; *p* = 1.0). The results were largely concordant with those obtained using the two other methods that we employed (Supporting Information Fig. S12–S14). The only discrepancy was in *Loranthus*, for which there was a correlation between the root-to-tip distances (*p* = 0.04); however, this comparison was based on few taxa (*n* = 13).

**Fig. 4.**
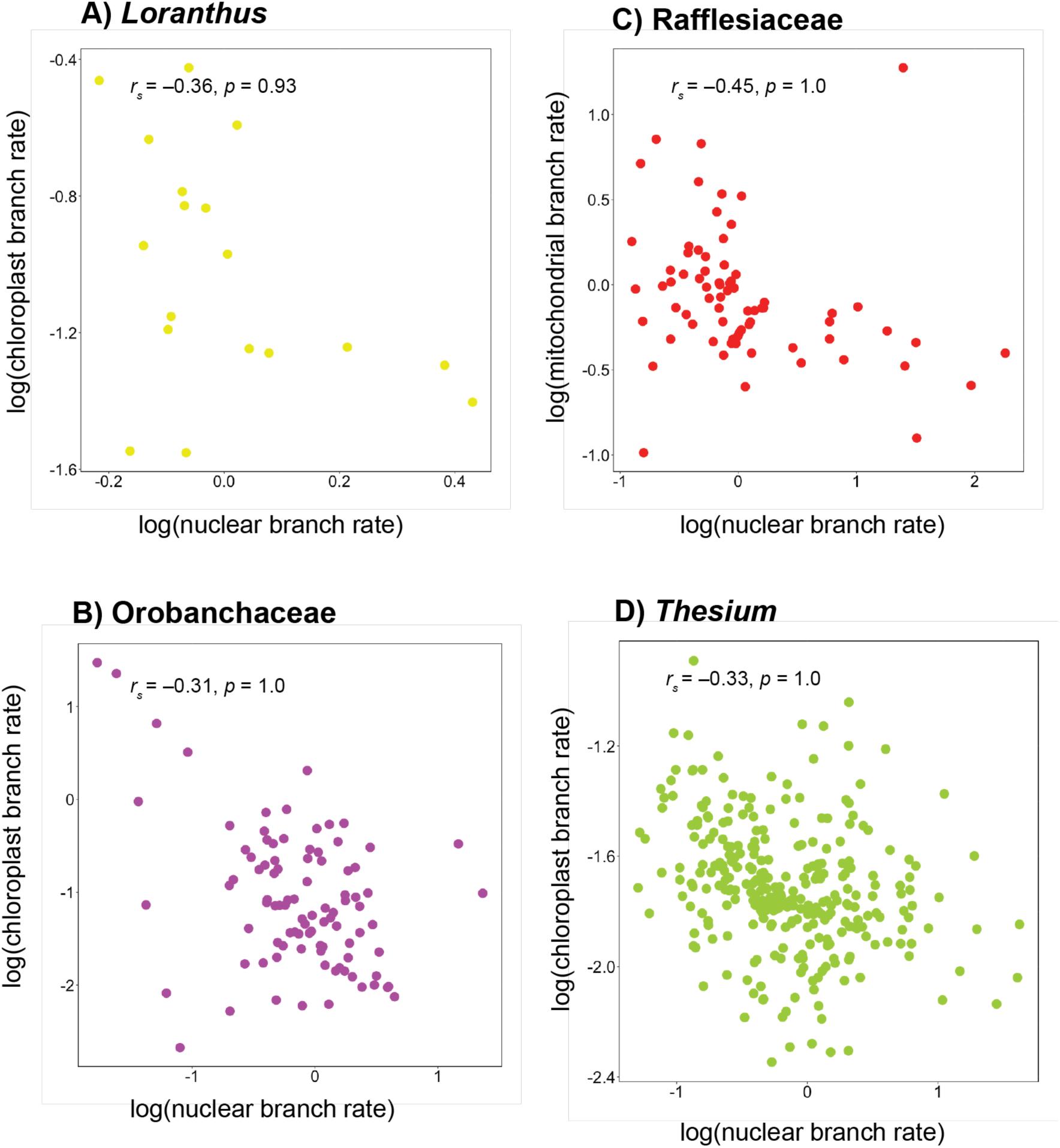
Rates of organellar evolution are decoupled in parasitic angiosperms. Comparisons of evolutionary branch rates of nuclear, mitochondrial, and chloroplast genes are shown for four parasitic lineages of angiosperms. Relative branch rates were estimated using Bayesian relaxed-clock analysis. The correlation coefficient (*r_s_*) and *p*-value (calculated using Spearman’s rank correlation test) are shown for each plot. Bold font indicates *p*-values below 0.05.

## Discussion

### Evolutionary rate correlations between genomes in land plants

Our study has shown that evolutionary rates are positively correlated between the nuclear, mitochondrial, and chloroplast genomes in the major lineages of land plants. These correlations were detected using various methods of analysis that we validated through an extensive simulation study. The pervasive signal of rate correlations across such diverse lineages is remarkable, given their ancient age (e.g., bryophyte mosses arose 424–416 Ma; Bechteler *et al.,* 2023) and their disparate life histories and ecologies. Contrastingly, we could not detect any evolutionary rate correlations in parasitic angiosperms. Indeed, parasitic angiosperms feature a variety of distinctive morphological and cellular traits that demarcate them from the rest of the flowering plants. For instance, the ‘parasite reduction syndrome’ phenomenon describes the convergent loss of vegetative tissue and photosynthetic organs and genes in parasitic flowering plants (Sanchez-Puerta *et al*., 2023).

Evolutionary rate correlations are consistent with the broad effects of cytonuclear covariation and interaction in land plants. These results are in line with the presence of mitonuclear covariation in animals (Weaver *et al*., 2022). There is evidence that coordination between the nuclear and organellar genomes in plants is vital to fundamental processes in plant cells, including ATP synthesis, oxidative phosphorylation, and photosynthesis (Sloan *et al*., 2014). For instance, in flowering plants, 2000 proteins are contained in the mitochondria, but fewer than 41 of these are encoded by the mitochondrial genome (Chen *et al*., 2017). The vast majority of proteins are encoded in the nucleus and transferred into the mitochondria by transit peptides. The nuclear genome can thus generate regulatory factors required for effective mitochondrial function (Chen *et al*., 2017; Hill, 2020; Wang *et al*., 2022). This is particularly important in the mitochondrial response to pathogen infection, activating programmed cell death, and immune function (Wang *et al*., 2022).

Gene exchange between genomic compartments might be partly driving the evolutionary rate correlation that we have uncovered. There are clear instances of bidirectional movement between the nuclear and mitochondrial genomes; for example, up to half of the maize mitochondrial genome is nuclear in origin (Chen *et al*., 2017). The movement of mitochondrial genes into the chloroplast genome appears to be much less profuse. The conserved chloroplast genome is more likely to transfer its genes to the nuclear and mitochondrial genomes, but unable to take up foreign organellar DNA (Chen *et al*., 2017; Hill, 2020). This is consistent with the uniformity in size and structure of the chloroplast genome in land plants. Furthermore, the sexually recombining nuclear genome has taken up hundreds of genes from the organellar compartments, as well as controlling and regulating many of the functions of the mitochondria and chloroplasts (Martin, 2003; Sloan *et al*., 2014). These features of genomic interaction suggest a weaker association between the evolutionary rates of mitochondrial and chloroplast genomes. However, if evolutionary rates in the mitochondrial and chloroplast genomes are both strongly linked to that of the nuclear genome, an indirect association can emerge between the rates of the two organellar genomes. This is consistent with the strength of the rate correlation being somewhat reduced between the two organellar compartments in ferns, angiosperms, and gymnosperms.

Our findings may also explain why higher rates of chloroplast substitution are linked to diversification in the angiosperm family Proteaceae (Duchêne & Bromham, 2013). A higher rate of change in the chloroplast genome predicts rapid changes in the nuclear and mitochondrial genomes, potentially driving adaptation, reproductive isolation, and eventually speciation. This is similar to findings in vertebrate and invertebrate animals, where mitonuclear covariation is a key factor in bolstering speciation (Hill, 2016; Wang *et al*., 2021; Weaver *et al*., 2022). Furthermore, a similar link between substitution rate and diversification has been found across many angiosperm clades (Lancaster, 2010; Bromham *et al*., 2015)

### Shared drivers of evolutionary rates

Although cytonuclear interactions are expected to produce some degree of evolutionary covariation between the nuclear and organellar genomes, the observed correlations in evolutionary rates might reflect a shared response by all three genomic compartments to some other factor. For example, the genomes might be responding to the same selection pressures or might have a shared system of DNA repair (Williams *et al*., 2019). We looked for signatures of these factors by performing separate tests of correlations for nonsynonymous and synonymous sites. We found correlations between the nonsynonymous rates of nuclear and organellar genomes, consistent with positive selection and genomic interactions driving the shared variation in evolutionary rates. However, we also found correlations between the synonymous rates of nuclear and organellar genomes, suggesting that there is a shared driver of mutation rates across the three genomic compartments. This is supported by the results of a previous study that found correlated rates at synonymous sites, but not at nonsynonymous sites (Bromham *et al*., 2015). However, our results suggest that the patterns of mutation rate variation are also being propagated to nonsynonymous rates, where a weaker correlation is apparent across the three genomes. The results of our analyses, conducted at a broad scale on hundreds of nuclear genes and tens of organellar genes, contradict those of previous studies that only found correlations between mitochondrial- and nuclear-encoded mitochondrial genes known to interact (Yan *et al*., 2019; Piccinini *et al*., 2021; Weaver *et al*., 2022).

Previous studies have identified a number of potential drivers of mutation rates in plant genomes. For instance, generation time is closely linked to long-term rates of meiosis in plants, since each elapsed generation has undergone a meiotic cycle. Thus, generation time might be driving the mutation rates of all three genomes (Petit & Hampe, 2006). Also, plant height is closely linked to generation time, and height has been found to be strongly associated with the rate of molecular evolution (Lanfear *et al*., 2013). This may be because taller plants have slower growth in their apical meristem and so undergo fewer cycles of mitosis (Li *et al*., 1996; Lanfear *et al*., 2013). Therefore, fewer mutations are inherited by offspring, effectively reducing the evolutionary rates of taller plants.

Environmental factors such as temperature can potentially act as mutagens in plants (Davies *et al*., 2004; Quiroz *et al*., 2023). This has been shown by experiments in *Arabidopsis thaliana*, where multi-generational exposure to higher temperature led to accumulation of mutations in genes involved in DNA repair, defense repair, and signalling, as well as increasing the mutation rate (Belfield *et al*., 2021; Lu *et al*., 2021). UV irradiation is another well-documented mutagen in plants (Davies *et al*., 2004; Quiroz *et al*., 2023), leading to oxidative DNA damage as demonstrated in Cucurbitaceae (Watanabe *et al*., 2006). These environmental drivers of mutation would potentially act on all three genomes in land plants.

Population size is also likely to have a broad impact on the evolutionary dynamics of nuclear, mitochondrial, and chloroplast genomes. There is less efficient selection against mutations in small populations, leading to a reduction in the efficiency of DNA repair machinery and ultimately to elevated mutation rates across the three genomes (Bergeron *et al*., 2023). There is evidence of unique systems of DNA repair in the nuclear, mitochondrial, and chloroplast genomes. For instance, mitochondria lack nucleotide excision repair, whereas chloroplasts are unable to fix double-strand breaks using non-homologous end joining (Boesch *et al*., 2011). Genome copy number and mutation rate are negatively linked in mitochondria, this may be because unmutated copies can perform repairs on double-strand breaks, however, this pattern is not seen in chloroplasts (Zwonitzer *et al*., 2024). Alternatively, there is mounting evidence for strong links between the epigenome and mutation rates in plants (Martincorena & Luscombe, 2013; Quiroz *et al*., 2023). Further work on DNA repair systems in plants is needed to clarify their role in mutation rate variation across nuclear and organellar genomes.

### Evolutionary rates in land plants

Across land plants, we found that nuclear genomes have evolved the most rapidly, followed by chloroplast genomes and mitochondrial genomes. These results are consistent with those of previous studies (Wolfe *et al*., 1987; Drouin *et al*., 2008; Smith, 2015). Overall, land plant nuclear genomes exhibit highly dynamic evolution, with the largest genomes 2400 times the size of the smallest; nuclear genomes also undergo rampant polyploidisation in land plants (Pellicer *et al*., 2018). Furthermore, nuclear genomes are responsible for most cell functions, encoding between 25,000–40,000 genes (Heslop-Harrison & Schmidt, 2012). Thus, their high evolutionary rate, when compared with the reduced rates of the chloroplast and mitochondrial genomes, is not unexpected. The low evolutionary rates of nuclear and organellar genomes in gymnosperms are consistent with the long generations of these taxa (Leitch & Leitch, 2012; De La Torre *et al*., 2017). Whilst many angiosperms can complete their life cycle in mere weeks, most gymnosperms take years to finish a single cycle.

There was a moderate amount of among-lineage rate heterogeneity overall in nuclear and chloroplast genes in the major land plant clades. Our results are consistent with previous findings of high-rate heterogeneity among lineages in angiosperm mitochondrial genomes. Whilst plants have drastically lower rates of mitochondrial evolution, the angiosperm mitochondrial genome is dynamic and exhibits highly variable evolutionary rates (Galtier, 2011; Havird *et al*., 2017; Petersen *et al*., 2020). The inflated rates of mitochondrial evolution in lineages such as *Plantago*, *Silene*, and *Viscum* have been attributed to genetic transfer from bacteria, viruses, and the nuclear and chloroplast genomes (Zervas *et al*., 2019). Indeed, the phenomenon of mitochondrial fusion is exhibited by *Amborella trichopoda*, the sister taxon to all other angiosperms (Rice *et al*., 2013). The mitochondrial genome of this species has fused with the whole genomes of three green algae and a moss species, and has received 82 copies of protein-coding and rRNA genes via horizontal transfer from other angiosperm species (Rice *et al*., 2013). Further insights will be gained from analysing larger amounts of data, particularly additional mitochondrial data; mitochondrial genomes are publicly available for only 142 species of plants, compared with 4445 chloroplast genomes (Petersen *et al*., 2020).

### Evolutionary rates in parasitic angiosperms

The parasitic flowering plants included in our study exhibited high levels of among-lineage rate heterogeneity in their nuclear and organellar genomes. Rate heterogeneity was especially pronounced in nuclear genes of Rafflesiaceae, in mitochondrial genes of *Loranthus*, and in chloroplast genes of Orobanchaceae. Given the extreme levels of rate variation, it is perhaps unsurprising that we could not detect any evolutionary rate correlations in the parasitic angiosperms that we tested: *Loranthus*, Orobanchaceae, Rafflesiaceae, and *Thesium*.

Parasitism has evolved at least 12 times across the angiosperms (Westwood *et al*., 2010; Nickrent, 2020), with an estimated 4750 species attacking the stems and roots of host plants for nutrition. Our data sets included Rafflesiaceae, a group of the most reduced holoparasites (Molina *et al*., 2014). Holoparasites do not perform any photosynthetic function and rely on their hosts for their nutritional requirements, with the most extreme form being endoparasitism. Indeed, Rafflesiaceae include endoparasites such as *Sapria*, which lack an independent body and grow directly on their host (Cai *et al*., 2021). Other parasite species that we analysed occur as hemiparasites, which have not lost photosynthetic function completely, and only partly rely on their host for heterotrophic nutrient demands (Těšitel *et al*., 2010). Hemiparasites can be either obligate, depending on their hosts for nutrients, or facultative, where they are able to live independently of their hosts (Těšitel *et al*., 2010). We analysed data from obligate parasites *Loranthus*, which often reside on the stems of their hosts (Liu *et al*., 2018), and *Thesium*, which includes both obligate and facultative herbaceous parasites that attach to the roots of their hosts. Orobanchaceae includes members that span the range of parasitic modes, from free-living hemiparasites to holoparasites that live on their hosts and leach water and nutrients directly from the hosts’ roots, e.g., in desert hyacinths (Nickrent, 2020; Thorogood *et al*., 2021).

The degree of parasitism seems to be linked to the amount of chloroplast reduction. For example, the holoparasitic members of Orobanchaceae have experienced widespread chloroplast gene losses, with members of Rafflesiaceae being the most extreme iteration of this gene loss, as they appear to have lost their chloroplast genomes entirely (Molina *et al*., 2014; Zervas *et al*., 2019). The hemiparasitic *Viscum*, a genus of mistletoe, have lost 10–22% of their chloroplast genes compared with their autotrophic relatives (Petersen *et al*., 2015; Zervas *et al*., 2019). These gene losses are likely to be associated also with the loss of signalling and regulatory genes in the nuclear and mitochondrial genomes. This may have caused the lack of evolutionary rate correlations between nuclear and organellar genomes in parasitic angiosperms, given the loss of nuclear-encoded genes.

Moreover, due to the intimate nature of the host–parasite association, horizontal gene transfer is rife, especially in the mitochondrial genomes of parasite species. In family Rafflesiaceae, it was found that 24–41% of mitochondrial genes were donated from the host plant (Xi *et al*., 2013). This provides another potential explanation for the lack of evolutionary rate correlations; the foreign genes have not undergone coordinated evolution with the native genes over millions of years.

## Conclusions

We have identified correlations in evolutionary rates across the nuclear, mitochondrial, and chloroplast genomes in land plants, but not in parasitic angiosperms. These associations suggest that the evolution of the three genomes is at least partly governed by a common factor that acts upon mutation rates. Further insight into these evolutionary dynamics can be gained through detailed studies of specific genes, including nuclear and mitochondrial genes that encode products that are used in the mitochondria. Our framework can be extended to other major lineages of plants, including the green algae but also red algae. We found a weakly positive (but insignificant) correlation in a small data set of chlorophytes. If there is evidence for evolutionary rate correlations in algae, this indicates that the dynamic is ancient and consistent across all Viridiplantae. Indeed, whether rate correlations exist at an even larger scale across between the genomic compartments across the Tree of Life is unknown. This is particularly pertinent as new organelles are being discovered, such as the nitrogen-fixing organelle, the ‘nitroplast’ (Coale *et al*., 2024). Another open question is whether evolutionary rate correlations would persist in other photosynthetic eukaryote lineages with plastids derived from secondary or tertiary symbioses, such as dinoflagellates (Maruyama & Kim, 2020). Uncovering the drivers of evolutionary rate correlations will allow better understanding of the causes of genomic change.

## Acknowledgements

This work was supported by the Australian Research Council (DP220103265). Y.A. also acknowledges the support provided by an Australian Government Research Training Program Scholarship. The authors acknowledge the Sydney Informatics Hub and the high-performance computing facility, Artemis, at the University of Sydney for providing computing resources. We also thank Samuli Lehtonen, Glenda Cardenas Ramirez, Miguel Angel García García, Pieter Pelser, Chi Toan Le, Daniel Nickrent, Sean Graham, and Frank E Anderson for providing access to data sets.

## Competing interests

There are no competing interests to report.

## Author contributions

YA, HS, and SYWH designed and planned the study. YA collected, curated, and analysed the empirical data sets and performed simulations. YA wrote the manuscript, with input from HS and SYWH.

## Notes

### Competing Interest Statement

The authors have declared no competing interest.

### Summary of Updates

Table 1 and Supplementary Information updated

