## Supplementary Information for "Evolutionary rates of nuclear and organellar genomes are linked in land plants"

The following Supporting Information is available for this article:

**Methods S1** Describes methods, results, and discussion of simulation study.

**Methods S2** Explains tests of the assumptions required for independent sister-pair contrasts.

**Notes S1** Brief discussion on detecting evolutionary rate correlations in relatively small data sets.

**Fig. S1** Performance of three methods for testing for correlations between nuclear and chloroplast evolutionary rates, for synthetic data generated with I) correlated and II) uncorrelated rates.

**Fig. S2** Boxplots depicting the proportion of replicates, for each method, where correlated rates of nuclear and chloroplast evolution were detected in analyses of synthetic data sets.

**Fig. S3** Boxplots summarising the proportion of replicates where correlated rates of nuclear and chloroplast evolution were detected in analyses of synthetic data sets under various settings, pooled across all three methods.

**Fig. S4** Boxplots summarising the proportion of replicates where correlated rates of nuclear and chloroplast evolution were detected in analyses of synthetic data sets under various settings, using root-to-tip distances.

**Fig. S5** Boxplots summarising the proportion of replicates where correlated rates of nuclear and chloroplast evolution were detected in analyses of synthetic data sets under various settings, using independent sister-pair contrasts.

**Fig. S6** Boxplots summarising the proportion of replicates where correlated rates of nuclear and chloroplast evolution were detected in analyses of synthetic data sets under various settings, using Bayesian branch rates.

**Fig. S7** Coefficient of variation of rates, a measure of among-lineage rate heterogeneity, estimated for each of the plant clades analysed in this study.

**Fig. S8** Rates of nuclear, mitochondrial, and chloroplast evolution are positively linked in the major embryophyte clades (using Bayesian branch rates).

**Fig. S9** Rates of nuclear, mitochondrial, and chloroplast evolution are positively linked in the major embryophyte clades (using root-to-tip distances).

**Fig. S10** Rates of nuclear, mitochondrial, and chloroplast evolution are positively linked in the major embryophyte clades (using independent sister-pair contrasts).

**Fig. S11** Rates of chloroplast and mitochondrial evolution are not significantly linked in Chlorophyta.

**Fig. S12** Rates of nuclear, mitochondrial, and chloroplast evolution are not linked in parasitic angiosperms (using Bayesian branch rates).

**Fig. S13** Rates of nuclear, mitochondrial, and chloroplast evolution are not linked in parasitic angiosperms (using root-to-tip distances).

**Fig. S14** Rates of nuclear, mitochondrial, and chloroplast evolution are not linked in parasitic angiosperms (using independent sister-pair contrasts).

**Table S1** Details of data sets analysed in this study.

**Table S2** Age of origin used for estimating absolute rates of substitution.

**Table S3** Coefficient of variation (CoV) of rates.

**Table S4** Results of tests of correlations between rates at $1^{st}$ + $2^{nd}$ codon sites and $3^{rd}$ codon sites, based on root-to-tip distances, for each of the organellar compartments in the major embryophyte clades.

**Table S5** Results of tests of correlations between rates at $1^{st}$ + $2^{nd}$ sites and $3^{rd}$ sites, using independent sister-pair contrasts, for each of the organellar compartments in the major embryophyte clades.

**Table S6** Evolutionary rates used to generate synthetic chloroplast and nuclear phylograms.

**Table S7** Parameters used to generate synthetic nuclear and chloroplast DNA sequences in *Seq-Gen*.

**Table S8** Details of independent sister-pair contrasts for nuclear and chloroplast genes, including assumption tests.

**Table S9** Details of independent sister-pair contrasts for nuclear and mitochondrial genes, including assumption tests.

**Table S10** Details of independent sister-pair contrasts for chloroplast and mitochondrial genes, including assumption tests.

**Table S11** Details of independent sister-pair contrast analyses for 1$^{st}$ + 2$^{nd}$ codon sites (‘nonsynonymous’ sites) in nuclear and chloroplast genes, including assumption tests.

**Table S12** Details of independent sister-pair contrasts for 3$^{rd}$ codon sites (‘synonymous’ sites) in nuclear and chloroplast genes, including assumption tests.

**Table S13** Details of independent sister-pair contrasts for 1$^{st}$ + 2$^{nd}$ codon sites (‘nonsynonymous’ sites) in nuclear and mitochondrial genes, including assumption tests.

**Table S14** Details of independent sister-pair contrasts for 3$^{rd}$ codon sites (‘synonymous’ sites) in nuclear and mitochondrial genes, including assumption tests.

**Table S15** Details of independent sister-pair contrasts for 1$^{st}$ + 2$^{nd}$ codon sites (‘nonsynonymous’ sites) in chloroplast and mitochondrial genes, including assumption tests.

**Table S16** Details of independent sister-pair contrasts for 3$^{rd}$ codon sites (‘synonymous’ sites) in chloroplast and mitochondrial genes, including assumption tests.

**Methods S1**

### Simulation Procedure

**Parameters used for generating synthetic data**

We carried out a simulation study to evaluate the performance of three methods for testing for correlations in evolutionary rates: i) root-to-tip distances, ii) independent sister-pair contrasts, and iii) correlations of branch rates inferred using Bayesian relaxed-clock analysis. The simulations were based on the maximum-clade-credibility chronogram of angiosperms reported by Magallón *et al.* (2015) and used by Sauquet *et al.* (2017). This dated tree had been inferred using Bayesian phylogenetic analysis of three chloroplast protein-coding genes (*atpB*, *rbcL*, and *matK*) and two nuclear genes (18S and 16S nuclear ribosomal DNA) from 792 angiosperm species. We simulated the evolution of 'nuclear' and 'chloroplast' sequences on three different trees that were obtained by pruning this reference tree according to a diversified sampling approach. These pruned trees had (*i*) 18 taxa, representing each of the major angiosperm lineages; (*ii*) 45 taxa, representing angiosperm orders; and (*iii*) 111 taxa, at the upper limit of computational feasibility for the simulations.

We obtained phylograms by multiplying the branch lengths of each of the three chronograms by a chosen mean evolutionary rate. The branch rates were generated using the R package *NELSI* (version 0.21) (Ho *et al.*, 2015). We used an uncorrelated lognormal relaxed clock (Drummond *et al.*, 2006), where the means of the branch-rate distributions were obtained from two different empirical estimates (low and high) of angiosperm nuclear and chloroplast evolution. Similarly, we generated three different levels of rate variation among lineages by setting the standard deviation of the lognormal distribution to 0.25 (low), 0.75 (moderate), and 1.25 (very high).

**Synthetic nuclear and chloroplast trees with correlated rates of evolution**

To generate nuclear and chloroplast data sets with correlated rates of evolution, we first used NELSI to generate a 'nuclear' phylogram, using an empirical estimate for the rate of evolution of nuclear genes, $9.646\times10^{-4}$ subs/site/Myr (Magallón *et al.*, 2015). We treated this as our 'low' mean rate of nuclear evolution. We obtained a comparatively 'high' mean rate of nuclear evolution, $1.447\times10^{-3}$ subs/site/Myr, from a phylogenomic study that analysed 1167 low-copy nuclear genes from 111 angiosperm species (Zhang *et al.*, 2020). This nuclear phylogram was then duplicated and its branch lengths multiplied by a scaling factor. The scaling factor was obtained by simply dividing the chloroplast rate of evolution by the nuclear rate of evolution. This procedure yields nuclear and chloroplast phylograms with proportional branch lengths, which is equivalent to a scenario in which their branch rates are correlated. All rates and scaling factors are given in Table S6. This scaling approach is similar to one that we used previously to generate synthetic molecular and morphological data sets (Asar *et al.*, 2023).

**Synthetic nuclear and chloroplast trees with uncorrelated rates of evolution**

We simulated the evolution of synthetic nuclear and chloroplast sequences with uncorrelated branch rates. This allowed us to assess the detection of false positives. To generate uncorrelated rates, we did not employ any scaling factor between nuclear and chloroplast phylograms, and instead simply generated the branch rates separately. We generated a nuclear phylogram using empirically estimated rates of evolution, with both a low mean rate ($9.646\times10^{-4}$ subs/site/Myr) and a high mean rate

($1.447\times10^{-3}$ subs/site/Myr), and a chloroplast phylogram similarly using an empirically estimated low mean rate ($5.957\times10^{-4}$ subs/site/Myr) and high mean rate ($8.417\times10^{-4}$ subs/site/Myr) (Table S6). The low rate of chloroplast evolution was inferred using 76 protein-coding genes from the chloroplast genomes of 193 angiosperm taxa by Foster *et al.* (2017). The high rate of chloroplast evolution was sourced from Magallón *et al.* (2015).

**Simulations of sequence evolution**

We performed simulations of DNA sequence evolution on the phylograms described above. These simulations were carried out using the GTR model in *Seq-Gen* (version 1.3.4) (Rambaut & Grassly, 1997) (Table S7), under a range of conditions as described above: three numbers of taxa, 18, 45, and 111; two rates of evolution for the nuclear and chloroplast data, with a high and a low rate for each; and three levels of rate variation among branches, low (0.25), moderate (0.75), and high (1.25). In addition, we generated data sets with two sequence lengths, 1000 and 10,000 bp. Collectively, these simulations involved a total of 36 different settings, and we generated 20 replicates for each of these settings for a total of 720 synthetic data sets each of alignments with correlated and uncorrelated evolutionary rates.

**Maximum-likelihood and Bayesian analysis**

To test for signals of correlated evolution, we used three methods: (i) root-to-tip distances, (ii) sister-pairs analysis, and (iii) correlations of branch rates inferred from Bayesian relaxed-clock analysis. The first two methods were performed on maximum-likelihood trees (phylograms) and the final method on the trees inferred using Bayesian phylogenetic analysis. For all analyses, the topology was constrained to match the tree used for simulation.

The maximum-likelihood analysis was performed in IQTREE2 (version 2.2.2.6) (Bui *et al.*, 2020), using the GTR+G+I model of nucleotide substitution. Following the estimation of branch lengths, we removed the gymnosperm outgroup, *Welwitschia mirabilis*. Bayesian phylogenetic analysis was performed using BEAST (versions 2.6.6 and 2.6.7) (Bouckaert *et al.*, 2019). The nuclear and chloroplast data were treated as distinct site partitions with unlinked clock models. The uncorrelated lognormal relaxed clock was used to model rate variation among branches (Drummond *et al.*, 2006). The Markov chain Monte Carlo (MCMC) analysis was run for 10,000,000 steps, sampling trees every 1000 steps. Two independent replicates were run and the samples were combined after the removal of the first 10% as burn-in. Following the maximum-likelihood and Bayesian phylogenetic analyses, we tested for correlations between the evolutionary rates of synthetic nuclear and chloroplast gene sequences, using the three methods mentioned above. These methods are described in detail in the main text.

### Simulation Results and Discussion

**Performance of methods**

Overall, the three methods that we evaluated in this study displayed high accuracy when applied to synthetic nuclear and chloroplast sequences, correctly detecting evolutionary rate correlations for 88.5% of replicates on average. The methods also had low detection of false positives, incorrectly detecting correlations in 3.8% of replicates on average (Fig. S1–S3). Analyses of branch rates inferred using Bayesian relaxed clocks were able to detect evolutionary rate correlations under a broad range of simulation settings (Fig. S1–S2, S6). Under this method, the mean correct detection of rate correlations was 99.7% across all settings. The detection of false positives was well below that expected by frequentist statistics, at only 2.1% across all replicates. Thus, we classed this as the most accurate (due to its low detection of false positives) and the most powerful method (due to its high detection of true rate correlations).

The Bayesian branch-rate method was closely followed by root-to-tip distances, which had an average detection of true correlations of 99.7%, but a slightly inflated detection of false positives, at 6.8% (Fig. S1–S2, S4). The method that performed most poorly was independent sister-pair contrasts (Fig. S1–S2, S5); as described in the main text, where the data set comprised only 18 taxa, the method was unable to detect evolutionary rate correlations because of the small number of contrasts sampled ($n$ = 3). Overall, independent sister-pair contrasts had an average detection of true rate correlations of 66.0%, but this increased to 99.0% when the 18-taxa setting was excluded. The detection of false positives was low, with an average detection of 2.5%, but this increased slightly to 3.8% when the 18-taxa setting was excluded. Our results confirm that independent sister-pair contrasts is the most conservative of the three methods tested, and demonstrate that the approach performs very poorly when the number of taxa sampled is small.

**Impacts of different simulation settings**

The overall performance of methods improved with increasing numbers of taxa in the data set. For instance, the accurate detection of evolutionary rate correlations across replicates increased from 66.1%, 99.3%, to 100% when there were 18, 45, and 111 taxa, respectively (Fig. S1, S3). Independent sister-pair contrasts had particularly poor detection of correlated evolutionary rates when the data set comprised only 18 taxa (0%). Increasing the number of taxa reduced the number of false positives for Bayesian branch rates, for instance, the average detection of false positives declined from 5.42% to 0.83% to 0% for 18, 45, and 111 taxa respectively. In contrast, the average detection of false positives for root-to-tip distances increased slightly when there were more taxa, from 4.17% to 7.50% to 8.75% when there were 18, 45, and 111 taxa, respectively. Thus, analysis of root-to-tip distances has a slightly inflated detection of false positives compared with the other two methods, particularly when the data set has at least 45 taxa. Increasing the sequence lengths of synthetic nuclear and chloroplast DNA from 1000 bp to 10,000 bp led to a slight improvement in the detection of true correlations, from 88.1% to 88.9% (Fig. S1, S3). The similarity of results between the analyses of short and long sequences might be due to the idealised properties of synthetic data and the matched models in our analyses. We expect that analyses of larger empirical data sets benefit more strongly than shown here from increased sequence length and gene sampling. Overall, increasing the size of the data set, either by increasing the number of taxa or the sequence length, had a predictable, positive impact on the detection of correlated evolutionary rates between synthetic nuclear and chloroplast sequences.

We also explored the effect of low and high evolutionary rates in synthetic nuclear and chloroplast sequences on detecting rate correlations. We found that varying the simulated rate of evolution had only a small impact, with correct detection of correlated rates being on average 88.2% and 88.7%, respectively. Similarly, there were only slight differences in correct detection under conditions of low (87.6%), moderate (88.9%), and high (88.9%) levels of among-lineage rate heterogeneity (Fig. S1, S3). Where among-lineage rate heterogeneity was extremely high for synthetic nuclear and chloroplast sequences, the detection of false positives was slightly higher, at 5.6%, compared with 2.9% and 2.9% for low and moderate levels of rate heterogeneity, respectively. This might be due to inflated noise and possible model misspecification induced by extreme degrees of rate heterogeneity. Nevertheless, the value of 5.6% is similar to the expected false-positive rate of 5% under frequentist statistics.

**Methods S2**

**Land Plant Genomes: Tests of the Assumptions for Independent Sister-Pair Contrasts**

For our analyses based on independent sister-pair contrasts, we carried out tests of the data to check for violations of the assumption of homogeneity of variances, as described by Garland *et al.* (1992), Freckleton (2000), and Welch and Waxman (2008). All contrasts were log-transformed for assumption tests. We began by checking that the standardised contrasts (absolute) were not associated with their standardisation factors ('Assumption Test 1'). The standardisation factor used was the square root of the sum of the branch lengths for the contrast. This test allowed us to identify cases with 'shallow divergences' causing poorly estimated branch lengths (Garland *et al.*, 1992; Welch & Waxman, 2008). We removed any 'shallow' contrasts that contributed to a significant trend in the assumption test. Where a trend remained even after unsuitable contrasts were removed, we used a two-sample sign test to test for correlations between contrasts, as recommended previously (Welch & Waxman, 2008).

We also checked that the mean branch-length sum of the contrasts was not associated with the absolute differences of the contrasts ('Assumption Test 2'). Any association would indicate that a non-parametric test or another transform should be used (Freckleton, 2000; Welch & Waxman, 2008). There were a few data points that violated Assumption Test 2, so all contrasts were analysed using non-parametric Spearman's rank correlation test. The results of the assumption tests are detailed in Tables S8–S16 for all comparisons.

**Notes S1**

**Brief Note on Preliminary Analyses**

We detected correlations in preliminary analyses of smaller data sets from land plants. For example, our analysis of Bayesian branch rates, using multilocus data from angiosperms, detected a correlation in evolutionary rates between two mitochondrial genes (*atp*A and *mat*R) and two chloroplast genes (*atp*B and *rbc*L) ($r_s$ = 0.24 and $p$ = 0.01, (Naumann *et al.*, 2013). Furthermore, our simulation study showed that correlations could be detected even when using sequence alignments comprising as few as 1000 nucleotides. Thus, given the accuracy and power of the methods that we evaluated, along with positive results obtained from smaller multi-locus data sets in land plants, we proceeded with testing for associations between nuclear and organellar rates of evolution in parasitic angiosperms even though the data sets were small in comparison with the 50+ loci in the data sets for major land plant lineages.

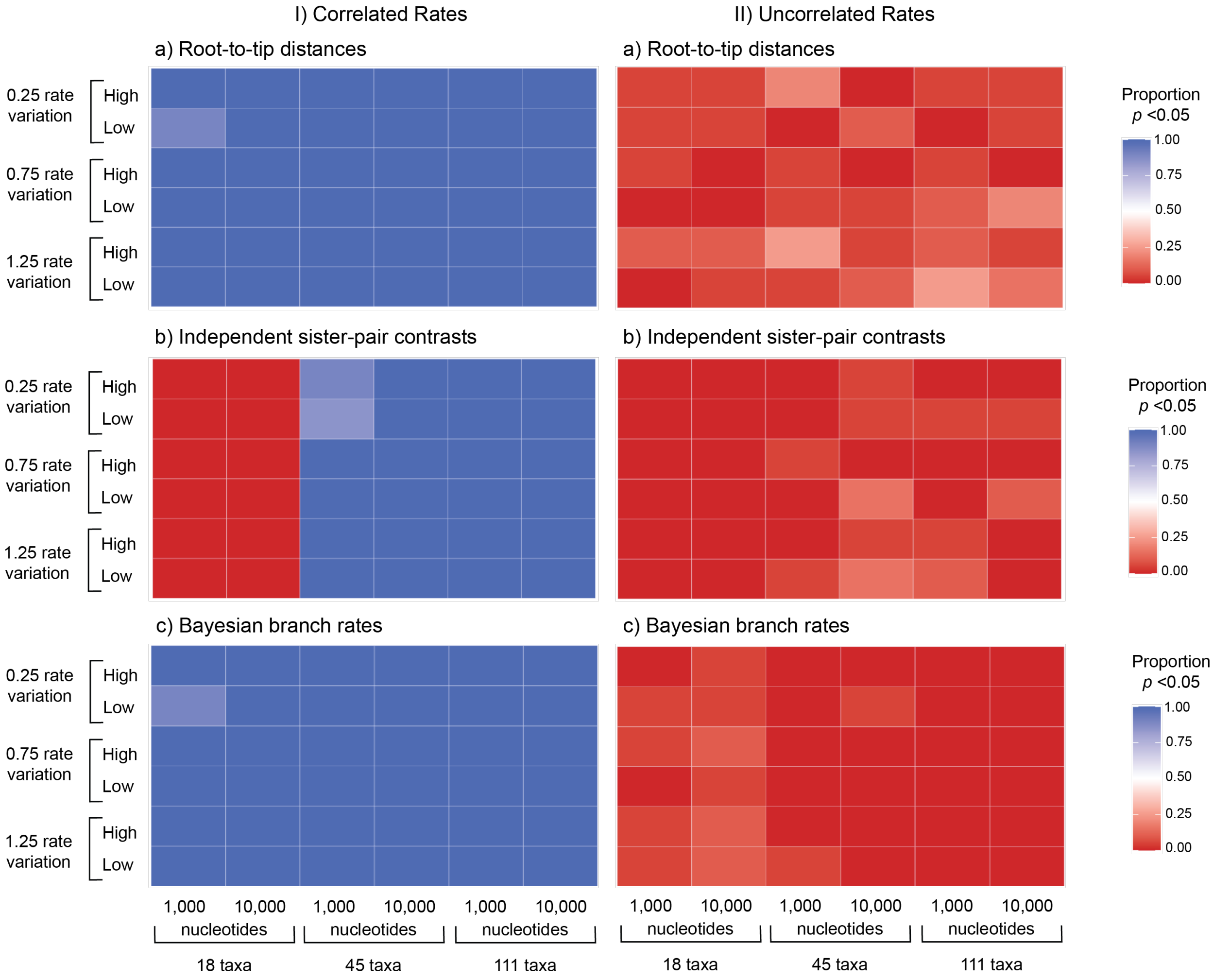


**Fig. S1. Performance of three methods for testing for correlations between nuclear and chloroplast evolutionary rates, for synthetic data generated with I) correlated and II) uncorrelated rates.** Heatmaps show results for analyses based on: a) root-to-tip distances, b) independent sister-pair contrasts, and c) correlations of branch rates inferred using Bayesian relaxed-clock analysis. Rows show results for three levels of among-lineage rate variation [0.25, 0.75, 1.25], and either low or high evolutionary rate. Columns show results for two sequence lengths [1000, 10,000] and three numbers of taxa [18, 45, 111]. Colors correspond to the proportion of 20 replicates for each setting that produced a significant positive correlation in evolutionary rates ($p < 0.05$).

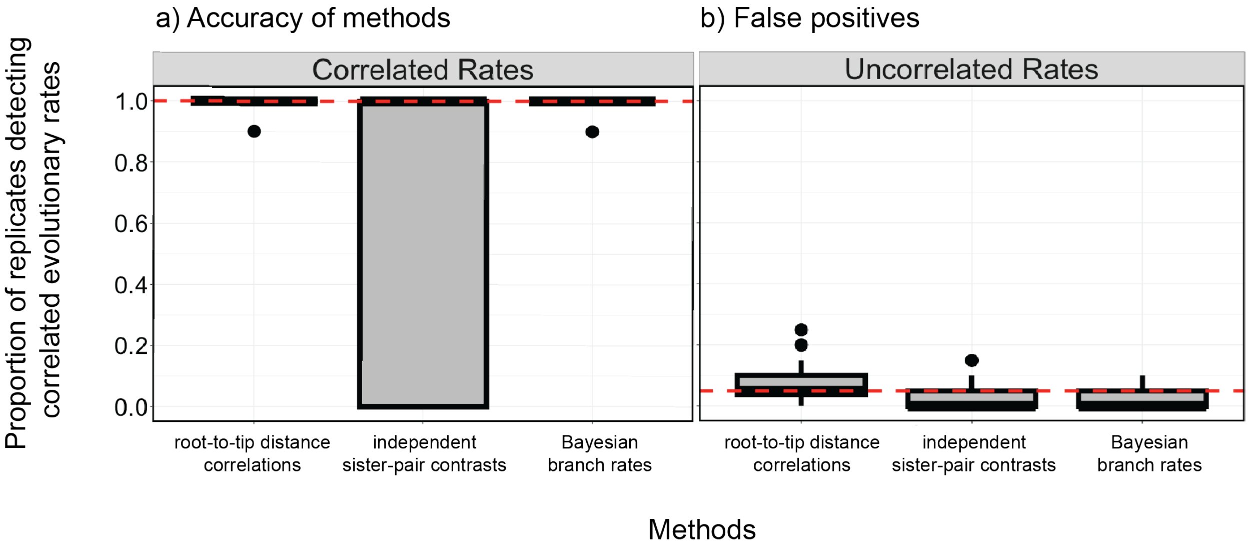


**Fig. S2. Boxplots depicting the proportion of replicates, for each method, where correlated rates of nuclear and chloroplast evolution were detected in analyses of synthetic data sets.** a) Accuracy of methods, measured as the proportion of replicates for which correlations between evolutionary rates were correctly detected. These data are sourced from synthetic nuclear and chloroplast sequences generated with correlated evolutionary rates. The dashed, horizontal red line highlights the ideal detection of correlations (100% of the time). b) False positives, shown as the proportion of replicates for which correlations between evolutionary rates were incorrectly detected. These data are sourced from synthetic nuclear and chloroplast sequences generated without correlated evolutionary rates. The dashed, horizontal red line highlights the false-positive detection expected under frequentist statistics (5% of the time).

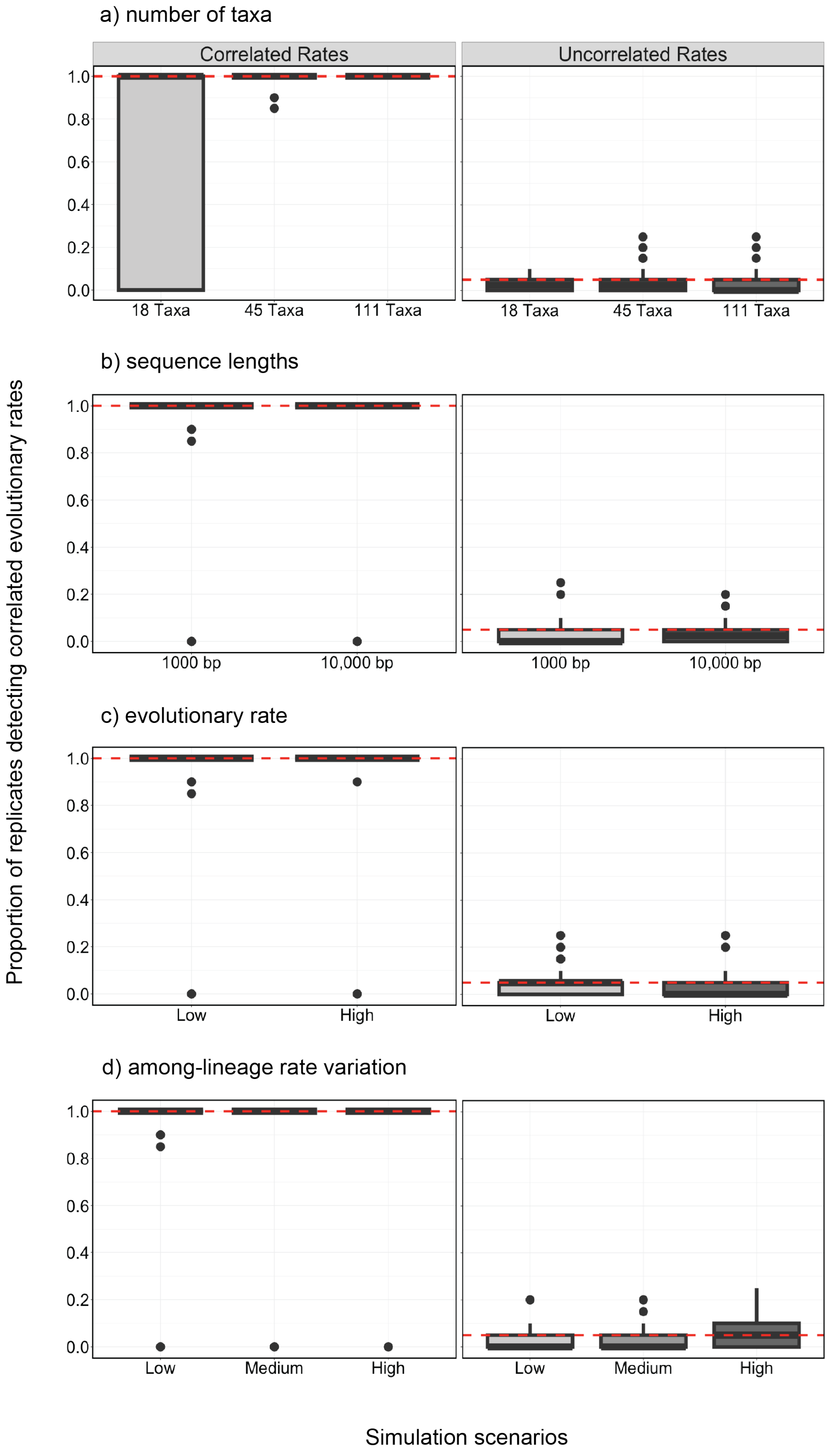

a) number of taxa
Correlated Rates
Uncorrelated Rates
1.0
0.8
0.6
0.4
0.2
0.0
18 Taxa
45 Taxa
111 Taxa
b) sequence lengths
1000 bp
10,000 bp
c) evolutionary rate
Low
High
d) among-lineage rate variation
Low
Medium
High
Proportion of replicates detecting correlated evolutionary rates
Simulation scenarios

**Fig. S3. Boxplots summarising the proportion of replicates where correlated rates of nuclear and chloroplast evolution were detected in analyses of synthetic data sets under various settings, pooled across all three methods.** The left column ‘Correlated Rates’ depicts the proportion of replicates for which rate correlations were correctly detected. These data are from synthetic nuclear and chloroplast sequences generated with correlated evolutionary rates. The dashed, horizontal red line highlights the ideal detection of correlations (100% of the time). The right column ‘Uncorrelated Rates’ depicts the proportion of replicates for which correlated rates were incorrectly detected. These data are from synthetic nuclear and chloroplast sequences generated without correlated evolutionary rates. The dashed, horizontal red line highlights the false-positive detection expected under frequentist statistics (5% of the time). The different settings used were: a) three tree sizes, b) two sequence alignment lengths, c) a low and high evolutionary rate for the synthetic nuclear and chloroplast sequences, and d) three levels of among-lineage rate variation.

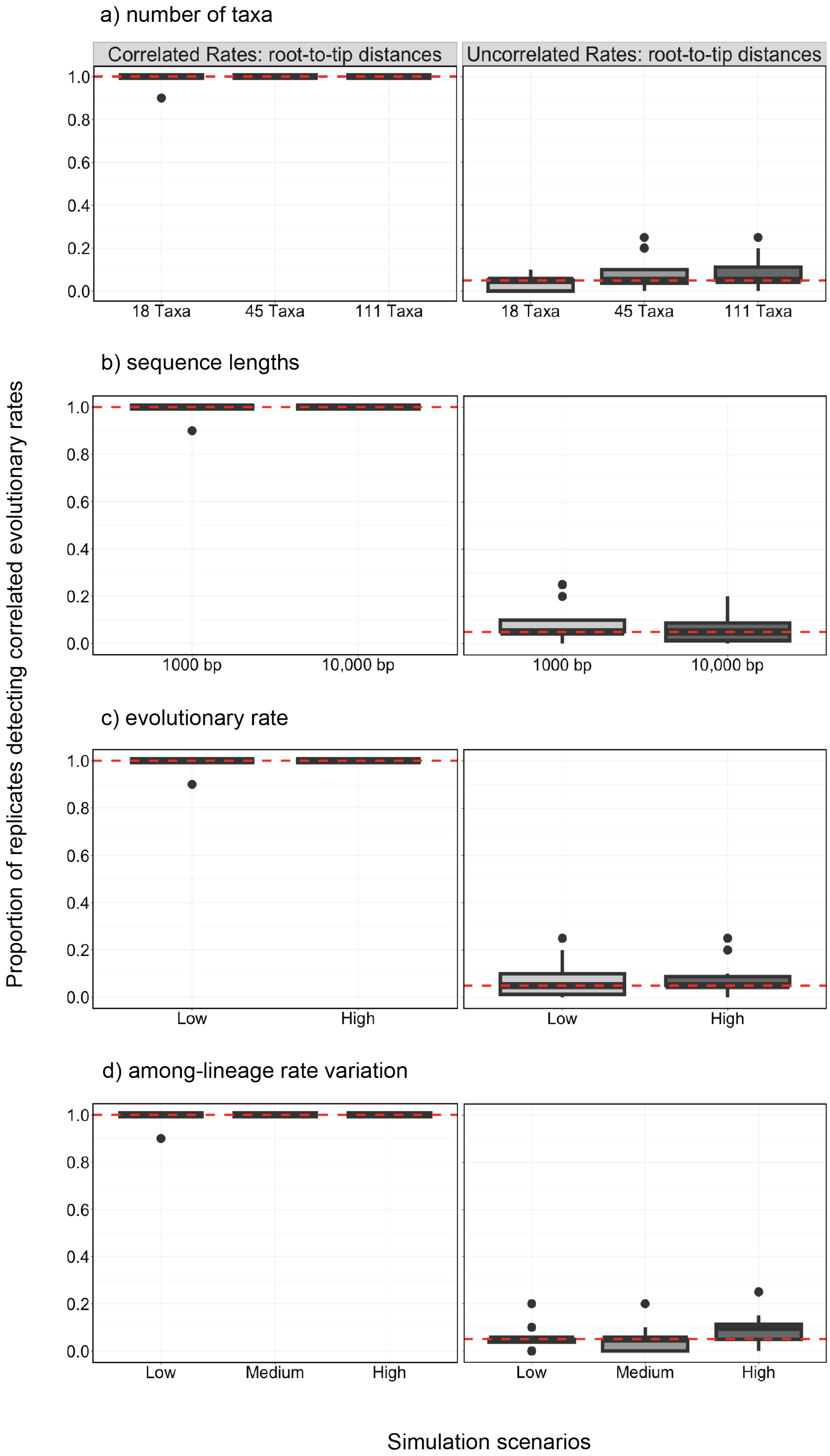
a) number of taxa
Correlated Rates: root-to-tip distances
Uncorrelated Rates: root-to-tip distances
1.0
0.8
0.6
0.4
0.2
0.0
18 Taxa
45 Taxa
111 Taxa
b) sequence lengths
1000 bp
10,000 bp
c) evolutionary rate
Low
High
d) among-lineage rate variation
Low
Medium
High
Proportion of replicates detecting correlated evolutionary rates
Simulation scenarios

215

**Fig. S4. Boxplots summarising the proportion of replicates where correlated rates of nuclear and chloroplast evolution were detected in analyses of synthetic data sets under various settings, using root-to-tip distances.** The left column 'Correlated Rates' depicts the proportion of replicates for which rate correlations were correctly detected. These data are from synthetic nuclear and chloroplast sequences generated with correlated evolutionary rates. The dashed, horizontal red line highlights the ideal detection of correlations (100% of the time). The right column 'Uncorrelated Rates' depicts the proportion of replicates for which rate correlations were incorrectly detected. These data are from synthetic nuclear and chloroplast sequences generated without correlated evolutionary rates. The dashed, horizontal red line highlights the false positive detection expected under frequentist statistics (5% of the time). The different settings used were: a) three tree sizes, b) two sequence alignment lengths, c) a low and high evolutionary rate for the synthetic nuclear and chloroplast sequences, and d) three levels of among-lineage rate variation.

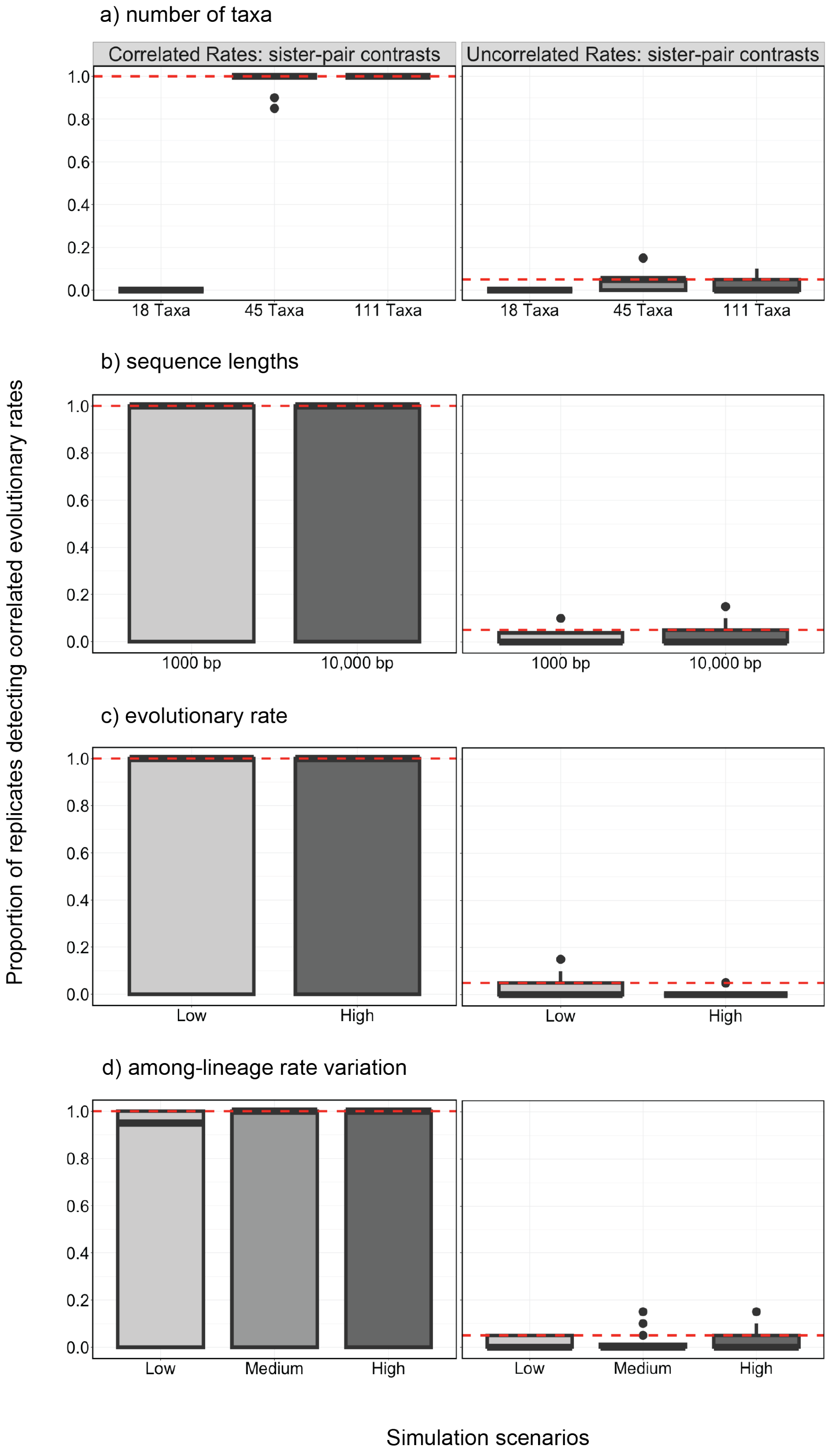
a) number of taxa
Correlated Rates: sister-pair contrasts
Uncorrelated Rates: sister-pair contrasts
1.0
0.8
0.6
0.4
0.2
0.0
18 Taxa
45 Taxa
111 Taxa
b) sequence lengths
1000 bp
10,000 bp
c) evolutionary rate
Low
High
d) among-lineage rate variation
Low
Medium
High
Proportion of replicates detecting correlated evolutionary rates
Simulation scenarios

229

**Fig. S5. Boxplots summarising the proportion of replicates where correlated rates of nuclear and chloroplast evolution were detected in analyses of synthetic data sets under various settings, using independent sister-pair contrasts.** The left column 'Correlated Rates' depicts the proportion of replicates for which rate correlations were correctly detected. These data are from synthetic nuclear and chloroplast sequences generated with correlated evolutionary rates. The dashed, horizontal red line highlights the ideal detection of correlations (100% of the time). The right column 'Uncorrelated Rates' depicts the proportion of replicates for which rate correlations were incorrectly detected. These data are from synthetic nuclear and chloroplast sequences generated without correlated evolutionary rates. The dashed, horizontal red line highlights the false-positive detection expected under frequentist statistics (5% of the time). The different settings used were: a) three tree sizes, b) two sequence alignment lengths, c) a low and high evolutionary rate for the synthetic nuclear and chloroplast sequences, and d) three levels of among-lineage rate variation.

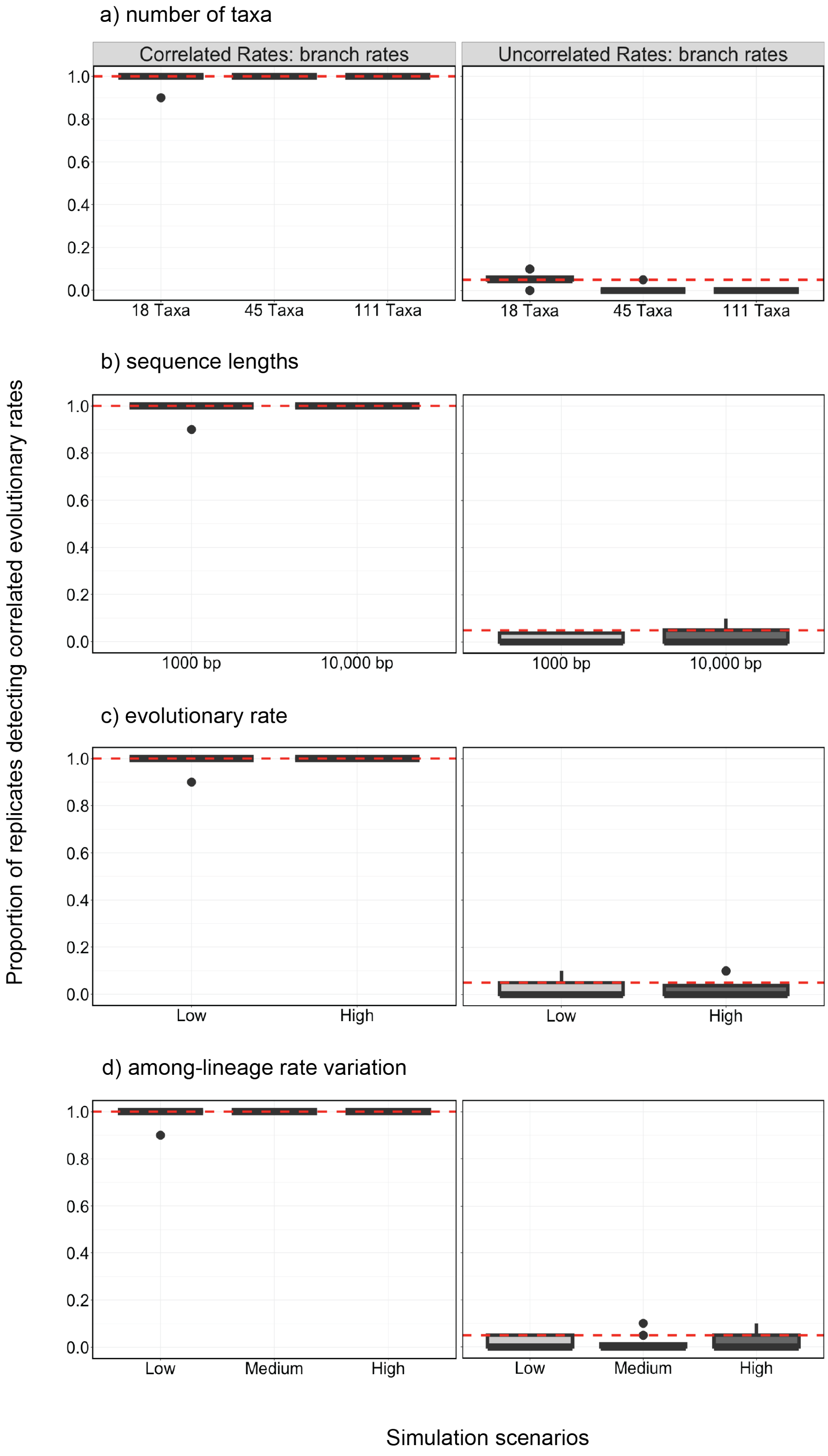

a) number of taxa
Correlated Rates: branch rates
Uncorrelated Rates: branch rates
1.0
0.8
0.6
0.4
0.2
0.0
18 Taxa
45 Taxa
111 Taxa
b) sequence lengths
1000 bp
10,000 bp
c) evolutionary rate
Low
High
d) among-lineage rate variation
Low
Medium
High
Proportion of replicates detecting correlated evolutionary rates
Simulation scenarios


243

**Fig. S6. Boxplots summarising the proportion of replicates where correlated rates of nuclear and chloroplast evolution were detected in analyses of synthetic data sets under various settings, using Bayesian branch rates.** The left column 'Correlated Rates' depicts the proportion of replicates for which rate correlations were correctly detected. These data are from synthetic nuclear and chloroplast sequences generated with correlated evolutionary rates. The dashed, horizontal red line highlights the ideal detection of correlations (100% of the time). The right column 'Uncorrelated Rates' depicts the proportion of replicates for which rate correlations were incorrectly detected. These data are from synthetic nuclear and chloroplast sequences generated without correlated evolutionary rates. The dashed, horizontal red line highlights the false-positive detection expected under frequentist statistics (5% of the time). The different settings used were: a) three tree sizes, b) two sequence alignment lengths, c) a low and high evolutionary rate for the synthetic nuclear and chloroplast sequences, and d) three levels of among-lineage rate variation.

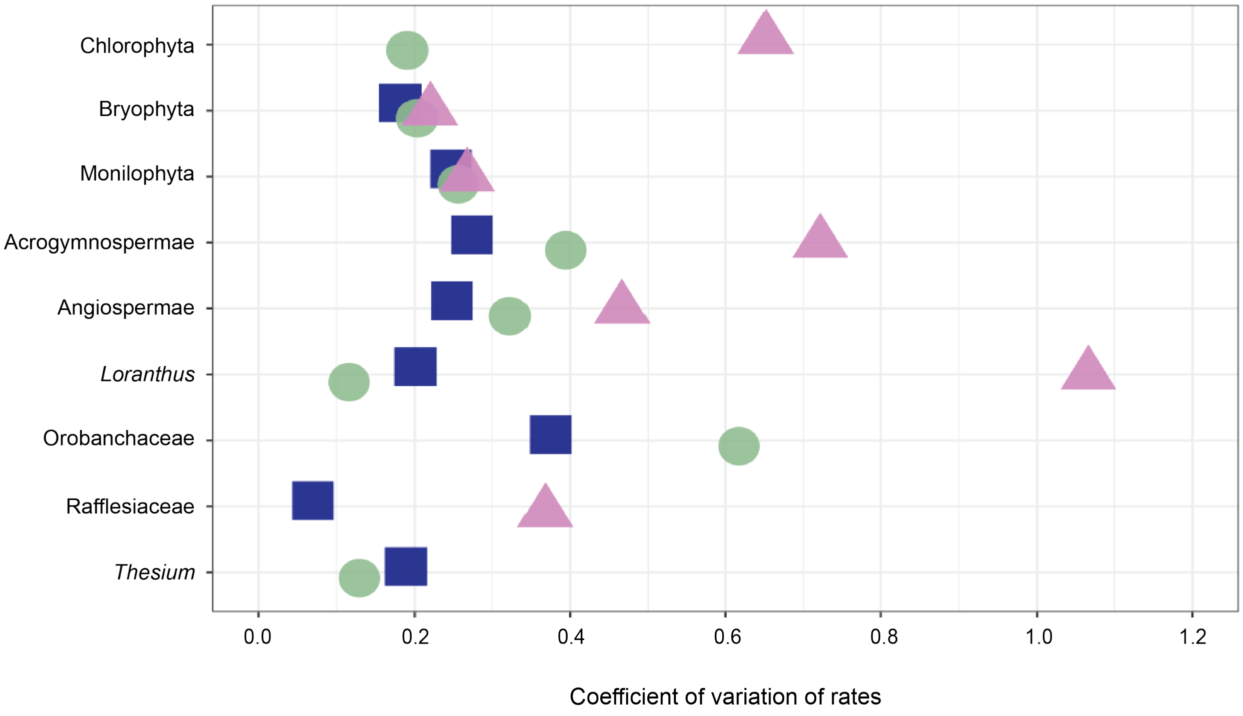


**Fig. S7. Coefficient of variation of rates, a measure of among-lineage rate heterogeneity, estimated for each of the plant clades analysed in this study**. The nuclear, mitochondrial, and chloroplast genomic compartments are represented by dark blue squares, violet triangles, and green circles, respectively. Estimates of the coefficient of variation were obtained by dividing the standard deviation of the root-to-tip distances by the mean root-to-tip distance. Root-to-tip distances were obtained from phylograms inferred using maximum likelihood.

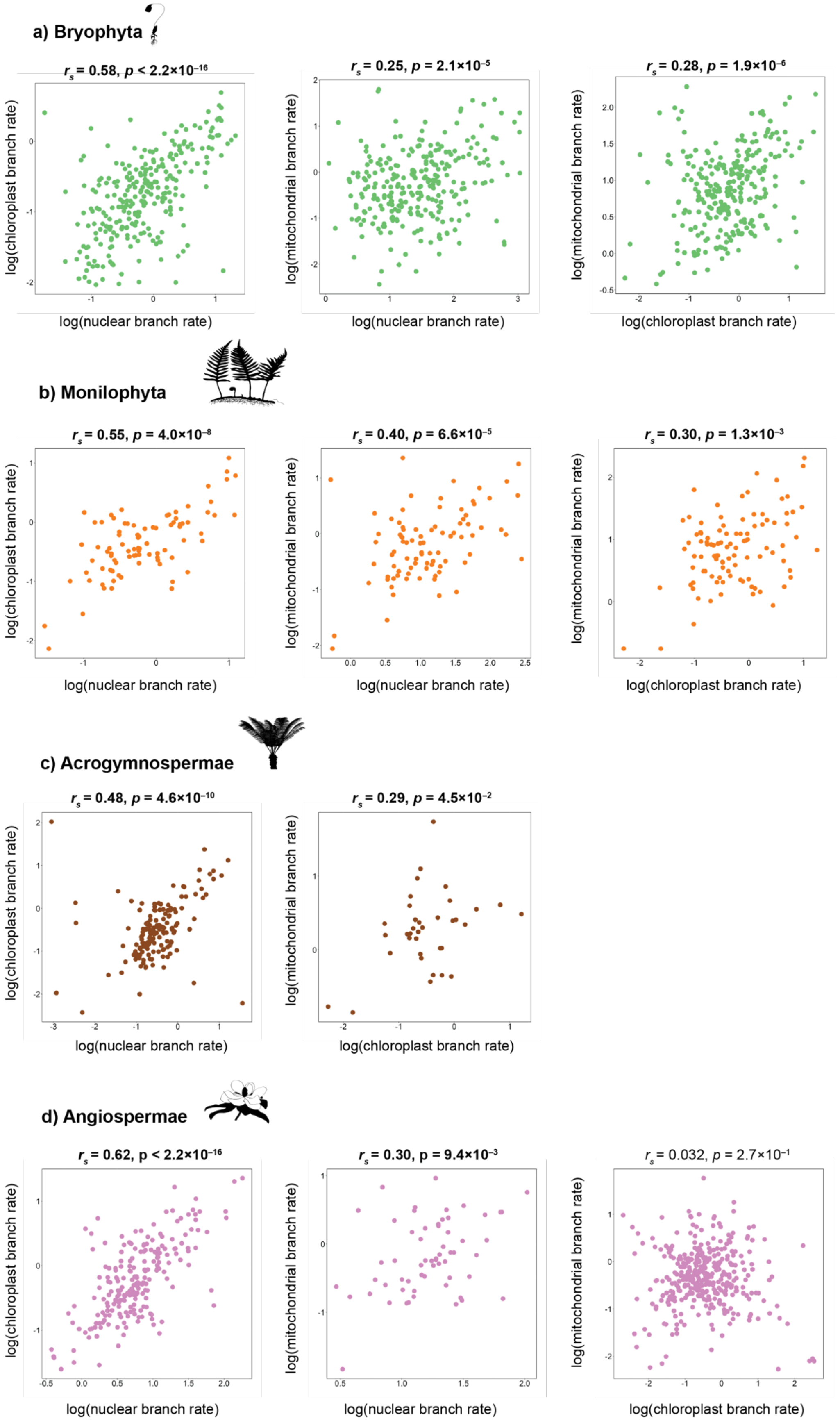
a) Bryophyta
$r_s$ = 0.58, $p < 2.2\times10^{-16}$
log(chloroplast branch rate)
log(nuclear branch rate)
$r_s$ = 0.25, $p = 2.1\times10^{-5}$
log(mitochondrial branch rate)
log(nuclear branch rate)
$r_s$ = 0.28, $p = 1.9\times10^{-6}$
log(mitochondrial branch rate)
log(chloroplast branch rate)
b) Monilophyta
$r_s$ = 0.55, $p = 4.0\times10^{-8}$
log(chloroplast branch rate)
log(nuclear branch rate)
$r_s$ = 0.40, $p = 6.6\times10^{-5}$
log(mitochondrial branch rate)
log(nuclear branch rate)
$r_s$ = 0.30, $p = 1.3\times10^{-3}$
log(mitochondrial branch rate)
log(chloroplast branch rate)
c) Acrogymnospermae
$r_s$ = 0.48, $p = 4.6\times10^{-10}$
log(chloroplast branch rate)
log(nuclear branch rate)
$r_s$ = 0.29, $p = 4.5\times10^{-2}$
log(mitochondrial branch rate)
log(chloroplast branch rate)
d) Angiospermae
$r_s$ = 0.62, p < $2.2\times10^{-16}$
log(chloroplast branch rate)
log(nuclear branch rate)
$r_s$ = 0.30, p = $9.4\times10^{-3}$
log(mitochondrial branch rate)
log(nuclear branch rate)
$r_s$ = 0.032, $p = 2.7\times10^{-1}$
log(mitochondrial branch rate)
log(chloroplast branch rate)

**Fig. S8. Rates of nuclear, mitochondrial, and chloroplast evolution are positively linked in the major embryophyte clades.** Log-transformed relative branch rates plotted for nuclear, mitochondrial, and chloroplast genomes for the major land plant clades. Branch rates were estimated using Bayesian relaxed-clock analysis. The correlation coefficient ($r_s$) and *p*-value (calculated using Spearman's rank correlation test) are shown for each plot. Bold font indicates *p*-values below 0.05. Plant silhouettes are in the public domain and available at http://www.phylopic.org.

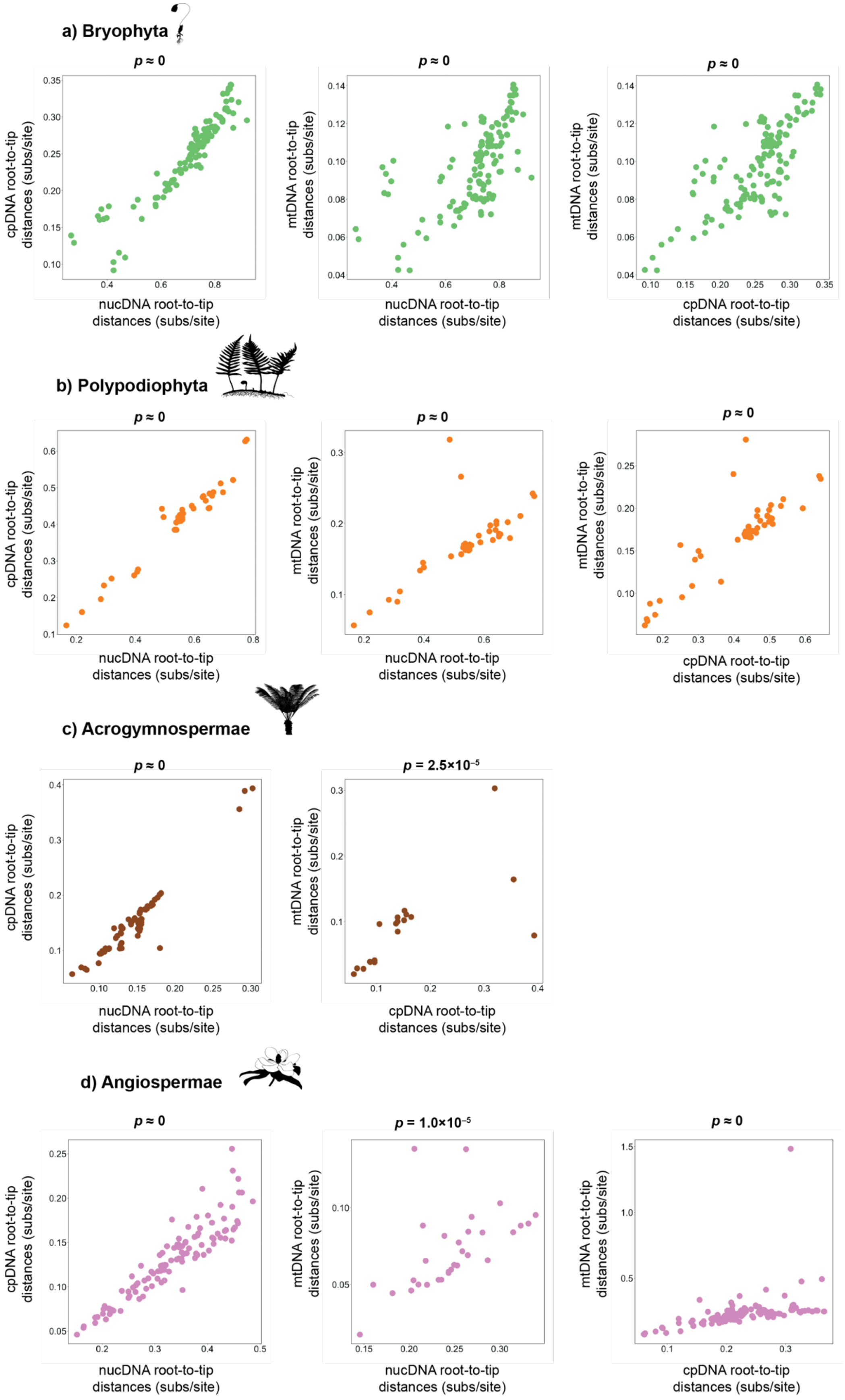
a) Bryophyta
b) Polypodiophyta
c) Acrogymnospermae
d) Angiospermae
p ≈ 0
p = 2.5×10⁻⁵
p = 1.0×10⁻⁵
cpDNA root-to-tip distances (subs/site)
mtDNA root-to-tip distances (subs/site)
nucDNA root-to-tip distances (subs/site)

**Fig. S9. Rates of nuclear, mitochondrial, and chloroplast evolution are positively linked in the major embryophyte clades.** Comparisons of root-to-tip distances plotted for nuclear, mitochondrial, and chloroplast genomes for the major land plant clades. Distances were calculated from phylograms inferred using maximum likelihood. The *p*-value (calculated using permutation correlation test) is shown for each plot. Bold font indicates *p*-values below 0.05. Plant silhouettes are in the public domain and available at http://www.phylopic.org.

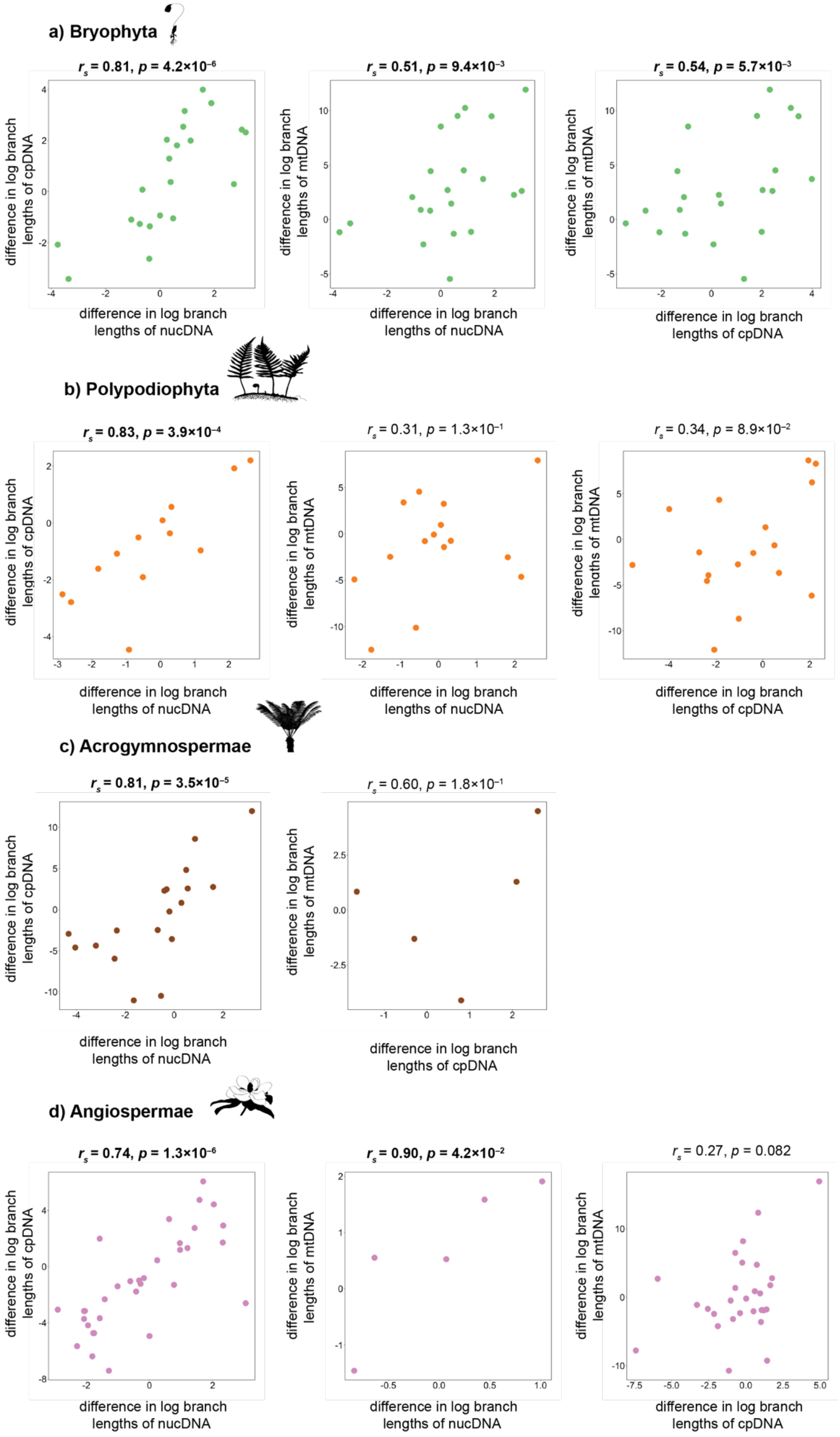
a) Bryophyta
$r_s$ = 0.81, $p$ = 4.2×10$^{-6}$
$r_s$ = 0.51, $p$ = 9.4×10$^{-3}$
$r_s$ = 0.54, $p$ = 5.7×10$^{-3}$
difference in log branch lengths of cpDNA
difference in log branch lengths of mtDNA
difference in log branch lengths of nucDNA
difference in log branch lengths of cpDNA
b) Polypodiophyta
$r_s$ = 0.83, $p$ = 3.9×10$^{-4}$
$r_s$ = 0.31, $p$ = 1.3×10$^{-1}$
$r_s$ = 0.34, $p$ = 8.9×10$^{-2}$
c) Acrogymnospermae
$r_s$ = 0.81, $p$ = 3.5×10$^{-5}$
$r_s$ = 0.60, $p$ = 1.8×10$^{-1}$
d) Angiospermae
$r_s$ = 0.74, $p$ = 1.3×10$^{-6}$
$r_s$ = 0.90, $p$ = 4.2×10$^{-2}$
$r_s$ = 0.27, $p$ = 0.082

**Fig. S10. Rates of nuclear, mitochondrial, and chloroplast evolution are positively linked in the major embryophyte clades.** Differences in log branch lengths between sister pairs plotted for nuclear, mitochondrial, and chloroplast genomes for the major land plant clades. Independent sister-pair contrasts were log-transformed and standardised. Contrasts were calculated from phylograms inferred using maximum likelihood. The correlation coefficient ($r_s$) and *p*-value (calculated using Spearman's rank correlation test) are shown for each plot. Bold font indicates *p*-values below 0.05. Plant silhouettes are in the public domain and available at http://www.phylopic.org.

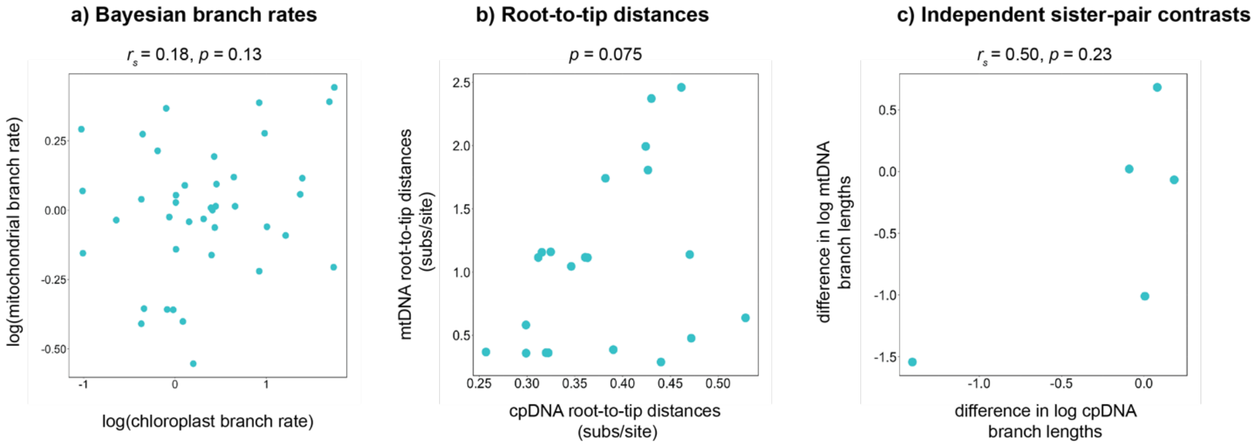


**Fig. S11. Rates of chloroplast and mitochondrial evolution are not significantly linked in Chlorophyta**. Results from three methods (a–c) used to test for evolutionary rate correlations between chloroplast and mitochondrial genes in Chlorophyta. The correlation coefficient ($r_s$) and *p*-value are shown for each plot.

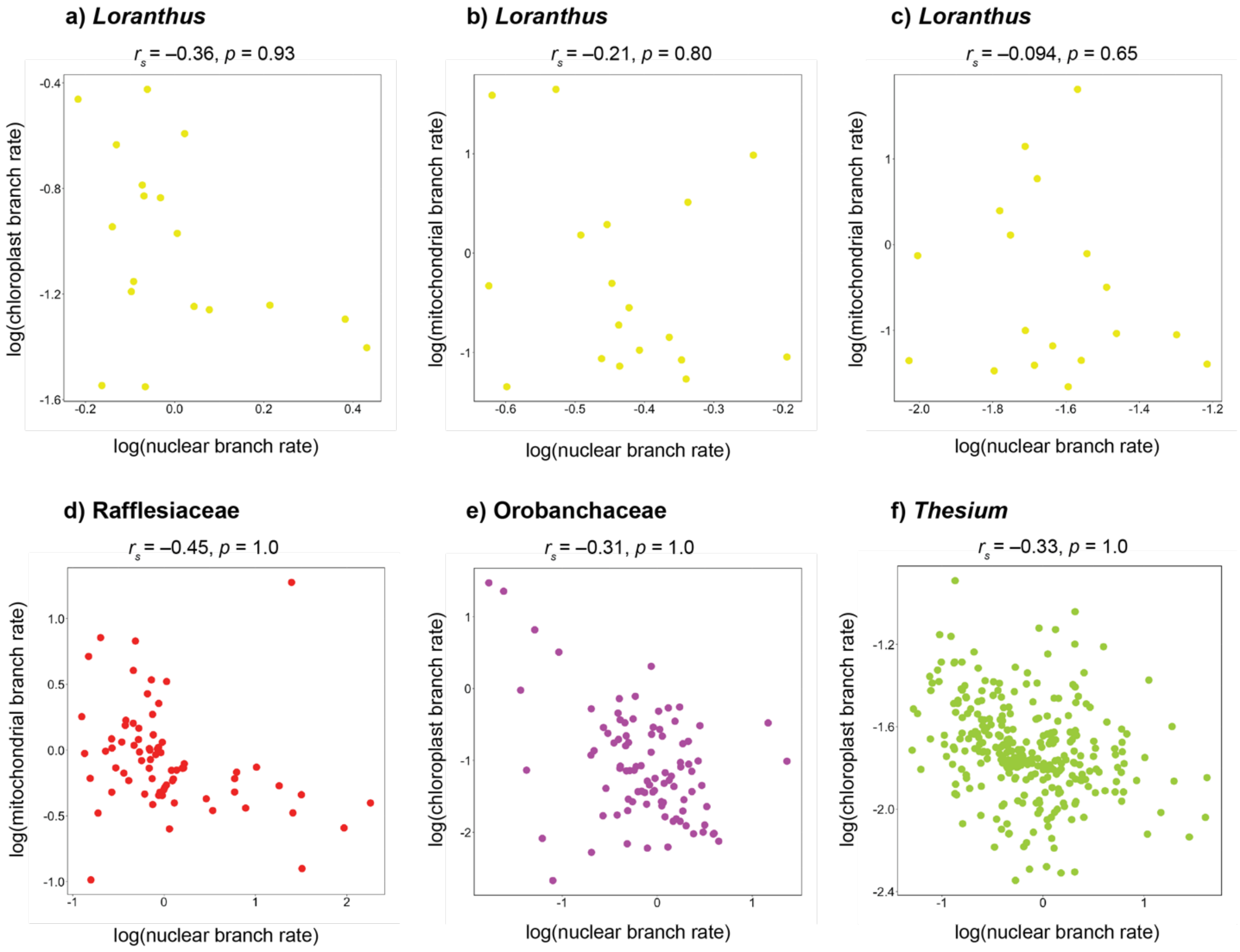


**Fig. S12. Rates of nuclear, mitochondrial, and chloroplast evolution are not linked in parasitic angiosperms.** Comparisons of log-transformed relative branch rates plotted for nuclear, mitochondrial, and chloroplast genes of parasitic flowering plants. Branch rates were estimated using Bayesian relaxed-clock analysis. The correlation coefficient ($r_s$) and *p*-value (calculated using Spearman's rank correlation test) are shown for each plot. Bold font indicates *p*-values below 0.05. Plant silhouettes are in the public domain and available at http://www.phylopic.org.

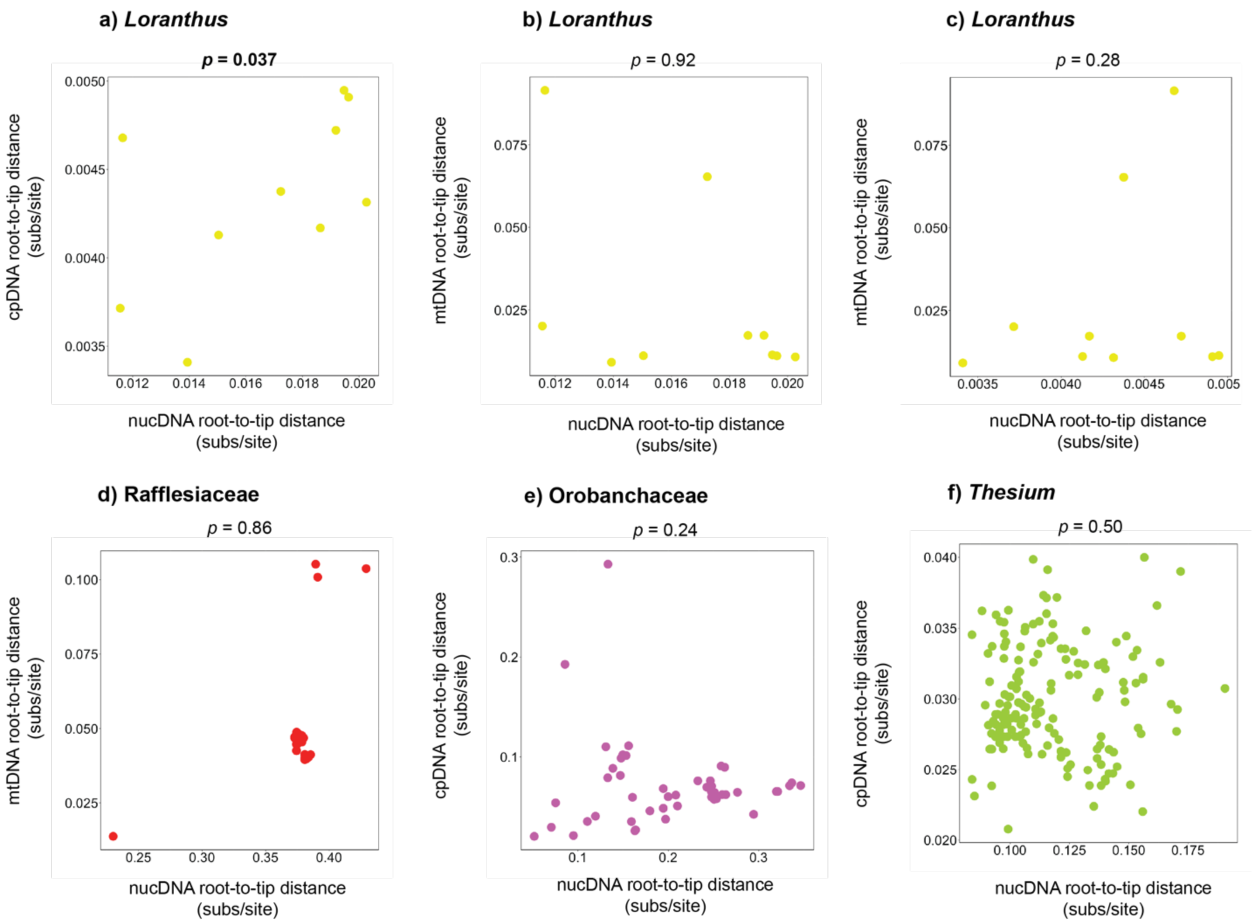


**Fig. S13. Rates of nuclear, mitochondrial, and chloroplast evolution are not linked in parasitic angiosperms.** Comparisons of root-to-tip distances plotted for nuclear, mitochondrial, and chloroplast genes of parasitic flowering plants. Root-to-tip distances were calculated using phylograms inferred using maximum likelihood. The *p*-value (calculated using permutation correlation test) is shown for each plot. Bold font indicates *p*-values below 0.05.

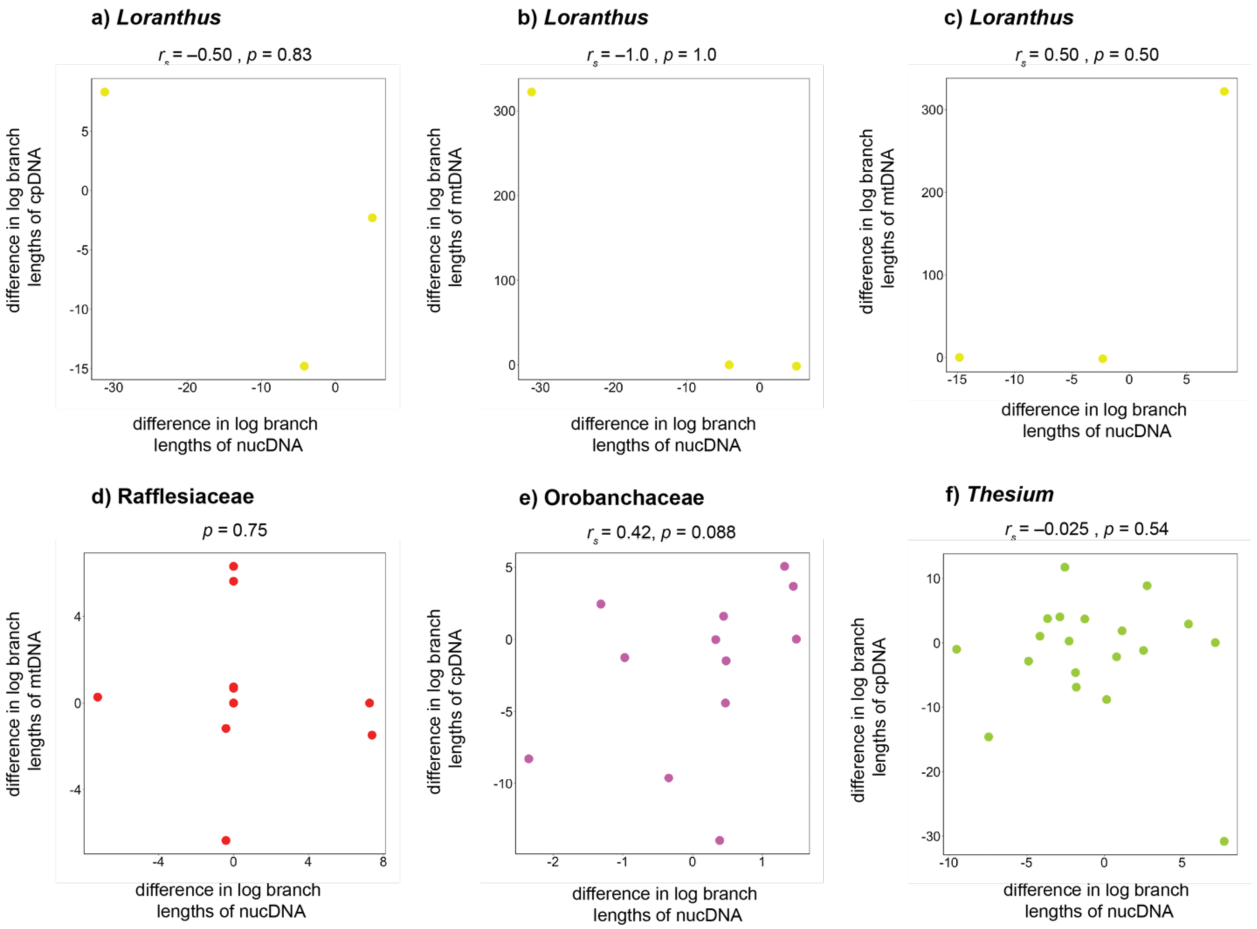


**Fig. S14. Rates of nuclear, mitochondrial, and chloroplast evolution are not linked in parasitic angiosperms.** Differences in log branch lengths between sister pairs plotted for nuclear, mitochondrial, and chloroplast genes of parasitic flowering plants. Independent sister-pair contrasts were log-transformed and standardised. Contrasts were calculated using phylograms inferred using maximum likelihood. The correlation coefficient ($r_s$) and *p*-value (calculated using Spearman's rank correlation test) are shown for each plot (excluding d) Rafflesiaceae, where no appropriate standardisation factor could be found, and a non-parametric two sample sign test was used). Bold font indicates *p*-values below 0.05.

**Table S1. Details of data sets analysed in this study.** Details are provided for sequence alignment length, number of genes, number of taxa, constraint topology, and the publication source of the sequence data.

| | Taxon | Nuclear Genes | Mitochondrial Genes | Chloroplast Genes | Taxa (in-group) | Constraint Topology[a] | Publication |
|---|---|---|---|---|---|---|---|
| (i) | Chlorophyta | - | 11,982 bp; 13 protein-coding genes | 30,803 bp; 45 protein-coding genes | 26 | Combined chloroplast and mitochondrial tree | Mekvipad and Satjarak (2019) |
| (ii) | Bryophyta | 154,548 bp; 105 protein-coding genes | 31,665 bp; 40 protein-coding genes | 64,368 bp; 82 protein-coding genes | 139 | Concatenated nuclear amino acid ML tree | Liu *et al.* (2019) |
| (iii) | Monilophyta | 35,877 bp; 25 low-copy loci[b] | - | - | 73 | Refer to (iv) | Rothfels *et al.* (2015) |
| (iv) | Monilophyta | - | 15,936 bp; 33 markers[b] | 26,854 bp; 31 markers[b] | 51 | Concatenated chloroplast marker tree | Cárdenas and Lehtonen (2023) |
| (v) | Acroymnospermae | 285,592 bp; 410 single-copy nuclear genes | - | - | 82 | Refer to (vi) | One Thousand Plant Transcriptomes Initiative (2019) |
| (vi) | Acroymnospermae | - | - | 57,015 bp; 78 genes | 890 | Dated chloroplast supermatrix tree | Stull *et al.* (2021) |
| (vii) | Acroymnospermae | - | 18,828 bp; 41 protein-coding genes | - | 20 | Refer to (vi) | Kan *et al.* (2021) |
| (viii) | Angiospermae | 285,592 bp; 410 single-copy genes | - | - | 837 | BEAST maximum-clade-credibility tree from Magallón *et al.* (2015)[c] | One Thousand Plant Transcriptomes Initiative (2019) |

| | | | | | | | |
|---|---|---|---|---|---|---|---|
| (ix) | Angiospermae | - | | 82,822 bp; 80 genes | 2,694 | Refer to (vii) | Li *et al.* (2019b) |
| (x) | Angiospermae | 76,833 bp; 59 low-copy genes | - | - | 60 | Refer to (x) | Zeng *et al.* (2014) |
| (xi) | Angiospermae | - | - | 58,347 bp; 78 protein-coding genes | 258 | Majority-rule bootstrap consensus tree using entire chloroplast supermatrix | Ruhfel *et al.* (2014) |
| (xii) | Angiospermae | - | 58,295 bp; 39 genes | 103,806 bp; 79 genes | 185 | Combined chloroplast and mitochondrial tree, dated | Tyszka *et al.* (2023) |
| (xiii) | *Loranthus* | 6,544 bp; Ribosomal DNA | 4,507 bp; 26S Ribosomal DNA | 148,114 bp; Plastome | 14 | Maximum-parsimony plastome tree | Nickrent *et al.* (2021) |
| (xiv) | Orobanchaceae | 5,085 bp; 6 markers (low-copy, ITS, and pentatricopeptide repeat) | - | 2,240 bp; 2 markers (*rps*2, *mat*K) | 54 | Maximum-likelihood tree using combined nuclear, chloroplast, and phytochrome loci | Li *et al.* (2019a) |
| (xv) | Rafflesiaceae | 814 bp; Ribosomal DNA, ITS | 6,662 bp; 3 markers (*atp*6, *mat*R, *nad*1 B-C intron) | - | 26 | Bayesian tree using combined nuclear and mitochondrial loci | Pelser *et al.* (2019) |
| (xvi) | *Thesium* | 748 bp; Ribosomal DNA, ITS | - | 3,336 bp; 2 markers (*trnDT*, *trnLF*) | 230 | Bayesian phylogram of the chloroplast tree | García *et al.* (2024) |

[a] The constraint topology was obtained from the same source publication, unless otherwise stated.
[b] Largely sourced from transcriptome data from the One Thousand Plant Transcriptomes Initiative (2019).
[c] The constraint topology used in the simulation study, used here to prune the taxon set to a manageable number.

**Table S2. Age of origin used for estimating absolute rates of substitution.** Point estimates from recent phylogenomic dating analyses used to infer the age of origin for Bryophyta, Monilophyta, Acrogymnospermae, and Angiospermae. Three estimates were used for Angiospermae, to reflect the uncertainty in their age of emergence; these estimates were sourced from the conservative, relaxed, and unconstrained analyses of Ramírez-Barahona *et al.* (2020).

| **Taxon** | **Age of Origin (Ma)** | **Publication** |
|---|---|---|
| Bryophyta | 419.49 | Bechteler *et al.* (2023) |
| Monilophyta | 423.2 | Nitta *et al.* (2022) |
| Acrogymnospermae | 325.37 | Ran *et al.* (2018) |
| Angiospermae[a] | 153.65 | Ramírez-Barahona *et al.* (2020) |
| Angiospermae[b] | 197.55 | Ramírez-Barahona *et al.* (2020) |
| Angiospermae[c] | 246.05 | Ramírez-Barahona *et al.* (2020) |

[a] Conservative calibrated estimate (complete) from Ramírez-Barahona *et al.* (2020)
[b] Relaxed calibrated estimate (complete) from Ramírez-Barahona *et al.* (2020)
[c] Unconstrained calibrated estimate (complete) from Ramírez-Barahona *et al.* (2020)

**Table S3. Coefficient of variation (CoV) of rates.** CoV estimated in a Bayesian framework (BEAST2) and using root-to-tip distances calculated from phylograms inferred using maximum likelihood. For the Bayesian estimates, the posterior mean CoV is reported, as well as the lower and upper limits of the 95% credibility interval (CI).

| Taxon | Genomic Compartment | Mean CoV | Lower 95% CI | Upper 95% CI | CoV root-to-tip distances |
|---|---|---|---|---|---|
| Chlorophyta | Chloroplast | 0.308 | 0.248 | 0.375 | 0.191 |
| | Mitochondrial | 0.903 | 0.784 | 1.04 | 0.652 |
| Bryophyta | Chloroplast | 0.702 | 0.639 | 0.777 | 0.204 |
| | Mitochondrial | 0.826 | 0.735 | 0.924 | 0.220 |
| | Nuclear | 0.784 | 0.703 | 0.860 | 0.182 |
| Monilophyta | Chloroplast | 0.712 | 0.606 | 0.839 | 0.256 |
| | Mitochondrial | 0.814 | 0.680 | 0.972 | 0.268 |
| | Nuclear | 0.743 | 0.654 | 0.844 | 0.247 |
| Acrogymnospermae | Chloroplast | 1.112 | 0.838 | 1.46 | 0.394 |
| | Mitochondrial | 0.906 | 0.704 | 1.13 | 0.722 |
| | Nuclear | 1.036 | 0.862 | 1.43 | 0.274 |
| Angiospermae | Chloroplast | 0.791 | 0.678 | 1.07 | 0.323 |
| | Mitochondrial | 1.52 | 1.35 | 1.71 | 0.467 |
| | Nuclear | 0.694 | 0.533 | 1.10 | 0.248 |
| *Loranthus* | Chloroplast | 0.439 | 0.258 | 0.608 | 0.116 |
| | Mitochondrial | 1.41 | 1.09 | 1.76 | 1.067 |
| | Nuclear | 0.276 | 0.046 | 0.538 | 0.202 |
| Orobanchaceae | Chloroplast | 1.406 | 0.961 | 1.92 | 0.618 |
| | Nuclear | 0.719 | 0.564 | 0.886 | 0.375 |
| Rafflesiaceae | Mitochondrial | 0.778 | 0.482 | 1.11 | 0.368 |
| | Nuclear | 1.60 | 0.702 | 2.59 | 0.0693 |
| *Thesium* | Chloroplast | 0.447 | 0.327 | 0.570 | 0.130 |
| | Nuclear | 0.871 | 0.686 | 1.07 | 0.189 |

**Table S4. Results of tests of correlations between rates at 1st + 2nd codon sites and 3rd codon sites, based on root-to-tip distances, for each of the organellar compartments in the major embryophyte clades.** For each organellar comparison, the *p*-value for a permutation correlation test and the number of in-group taxa sampled (*n*) are reported. Bold font indicates *p*-values below 0.05.

| Taxon | | Nuclear–Chloroplast | | Nuclear–Mitochondrial | | Chloroplast–Mitochondrial | |
|---|---|---|---|---|---|---|---|
| | | 1st + 2nd | 3rd | 1st + 2nd | 3rd | 1st + 2nd | 3rd |
| Bryophyta | *p* | **≈0** | **≈0** | **≈0** | **≈0** | **≈0** | **≈0** |
| | *n* | 135 | | 135 | | 135 | |
| Monilophyta | *p* | **≈0** | **≈0** | **≈0** | **≈0** | **≈0** | **≈0** |
| | *n* | 44 | | 44 | | 51 | |
| Angiospermae | *p* | **≈0** | **≈0** | - | - | **≈0** | **≈0** |
| | *n* | 30 | | - | | 185 | |

**Table S5. Results of tests of correlations between rates at 1st + 2nd sites and 3rd sites, using independent sister-pair contrasts, for each of the organellar compartments in the major embryophyte clades.** For each organellar comparison, the correlation coefficient ($r_s$) and $p$-value for a Spearman rank correlation test are reported, as well as the number of independent contrasts sampled ($n$) in the test. Bold font indicates $p$-values below 0.05.

| Taxon | | Nuclear–Chloroplast | | Nuclear–Mitochondrial | | Chloroplast–Mitochondrial | |
|---|---|---|---|---|---|---|---|
| | | **1st + 2nd** | **3rd** | **1st + 2nd** | **3rd** | **1st + 2nd** | **3rd** |
| Bryophyta | $r_s$ | 0.58 | 0.74 | 0.35 | 0.62 | 0.50 | 0.58 |
| | $p$ | $\mathbf{1.4{\times}10^{-4}}$ | $\mathbf{6.8{\times}10^{-7}}$ | $7.2{\times}10^{-2}$ | $\mathbf{2.1{\times}10^{-3}}$ | $\mathbf{1.5{\times}10^{-2}}$ | $\mathbf{1.4{\times}10^{-2}}$ |
| | $n$ | 36 | 35 | 19 | 20 | 19 | 15 |
| Monilophyta | $r_s$ | 0.49 | 0.8 | 0.53 | 0.27 | 0.27 | 0.13 |
| | $p$ | $\mathbf{4.4{\times}10^{-2}}$ | $\mathbf{2.6{\times}10^{-3}}$ | $7.4{\times}10^{-2}$ | $2.0{\times}10^{-1}$ | $1.4{\times}10^{-1}$ | $3.3{\times}10^{-1}$ |
| | $n$ | 13 | 11 | 9 | 12 | 17 | 15 |
| Angiospermae | $r_s$ | 0.79 | 0.64 | - | - | 0.29 | 0.16 |
| | $p$ | $\mathbf{2.4{\times}10^{-2}}$ | $6.9{\times}10^{-2}$ | - | - | $8.2{\times}10^{-2}$ | $2.1{\times}10^{-1}$ |
| | $n$ | 7 | 7 | - | - | 25 | 30 |

**Table S6. Evolutionary rates used to generate synthetic chloroplast and nuclear phylograms.**

| Evolutionary Rate | Nuclear Rate (subs/site/Myr) | Chloroplast Rate (subs/site/Myr) | Empirical Scaling Factor |
|---|---|---|---|
| High | $1.447\times10^{-3}$ | $8.417\times10^{-4}$ | 0.5817 |
| Low | $9.646\times10^{-4}$ | $5.957\times10^{-4}$ | 0.6176 |

**Table S7. Parameters used to generate synthetic nuclear and chloroplast DNA sequences in *Seq-Gen*.** These parameters were estimated by Sauquet *et al.* (2017) using data from Magallón *et al.* (2015).

| Parameter | Nuclear | Chloroplast |
|---|---|---|
| α shape of the gamma distribution | 0.525 | 0.666 |
| Gamma categories | 4 | 4 |
| Proportion of invariant sites | 0.274 | 0.188 |
| Pairwise transition rates (A↔C, A↔G, A↔T, C↔G, C↔T, G↔T) | 0.125, 0.35, 0.254, 0.06317, 1, 0.121 | 0.429, 0.8, 0.118, 0.28, 1, 0.306 |
| Nucleotide frequencies (A, C, G, T) | 0.241, 0.237, 0.299, 0.223 | 0.290, 0.187, 0.214, 0.309 |

**Table S8. Details of independent sister-pair contrasts for nuclear and chloroplast genes, including assumption tests.** Third column reports the number of independent contrasts used in analyses after removing inappropriate contrasts. Fourth to seventh columns give Kendall's tau ($\tau$) and *p*-values for both assumption tests. Any trend in the assumption tests is indicated by bold font ($p < 0.05$).

| Taxon | Data Type | Number of Contrasts | Assumption Test 1 | | Assumption Test 2 | |
|---|---|---|---|---|---|---|
| | | | Kendall's $\tau$ | *p*-value | Kendall's $\tau$ | *p*-value |
| Monilophyta | Chloroplast | 13 | –0.282 | 0.204 | –0.0256 | 0.952 |
| | Nuclear | | –0.103 | 0.675 | 0.179 | 0.435 |
| Bryophyta | Chloroplast | 21 | –0.229 | 0.158 | –0.229 | 0.158 |
| | Nuclear | | –0.210 | 0.197 | –0.105 | 0.531 |
| Acrogymnospermae | Chloroplast | 18 | –0.190 | 0.294 | 0.00654 | 1.0 |
| | Nuclear | | –0.176 | 0.330 | –0.0327 | 0.881 |
| Angiospermae | Chloroplast | 33 | –0.0795 | 0.528 | 0.208 | 0.0914 |
| | Nuclear | | 0.0341 | 0.794 | 0.242 | **0.0485** |
| *Loranthus* | Chloroplast | 3 | –0.333 | 1.0 | –0.333 | 1.0 |
| | Nuclear | | –0.333 | 1.0 | –0.333 | 1.0 |
| Orobanchaceae | Chloroplast | 12 | –0.182 | 0.459 | 0.0303 | 0.945 |
| | Nuclear | | 0 | 1.0 | 0.0909 | 0.737 |
| *Thesium* | Chloroplast | 19 | –0.123 | 0.489 | –0.146 | 0.406 |
| | Nuclear | | –0.170 | 0.332 | –0.146 | 0.406 |

**Table S9. Details of independent sister-pair contrasts for nuclear and mitochondrial genes, including assumption tests.** Third column reports the number of independent contrasts used in analyses after removing inappropriate contrasts. Fourth to seventh columns give Kendall's tau ($\tau$) and *p*-values for both assumption tests. Any trend in the assumption tests is indicated by bold font ($p < 0.05$). Note that we used a non-parametric sign-test for comparisons in Rafflesiaceae, as both assumption tests were severely violated even after removal of inappropriate contrasts and standardisation.

| Taxon | Data Type | Number of Contrasts | Assumption Test 1 | | Assumption Test 2 | |
|---|---|---|---|---|---|---|
| | | | Kendall's $\tau$ | *p*-value | Kendall's $\tau$ | *p*-value |
| Monilophyta | Mitochondrial | 15 | –0.162 | 0.435 | 0.162 | 0.435 |
| | Nuclear | | –0.143 | 0.495 | 0.105 | 0.627 |
| Bryophyta | Mitochondrial | 21 | –0.238 | 0.140 | –0.152 | 0.354 |
| | Nuclear | | –0.210 | 0.197 | –0.105 | 0.531 |
| Angiospermae | Mitochondrial | 5 | 0 | 1.0 | 0.2 | 0.817 |
| | Nuclear | | –0.6 | 0.233 | –0.4 | 0.483 |
| *Loranthus* | Mitochondrial | 3 | –0.333 | 1.0 | –0.333 | 1.0 |
| | Nuclear | | –0.333 | 1.0 | –0.333 | 1.0 |

**Table S10. Details of independent sister-pair contrasts for chloroplast and mitochondrial genes, including assumption tests.** Third column reports the number of independent contrasts used in analyses after removing inappropriate contrasts. Fourth to seventh columns give Kendall's tau ($\tau$) and *p*-values for both assumption tests. Any trend in the assumption tests is indicated by bold font ($p < 0.05$).

| Taxon | Data Type | Number of Contrasts | Assumption Test 1 | | Assumption Test 2 | |
|---|---|---|---|---|---|---|
| | | | Kendall's $\tau$ | *p*-value | Kendall's $\tau$ | *p*-value |
| Monilophyta | Chloroplast | 17 | –0.147 | 0.440 | 0.206 | 0.271 |
| | Mitochondrial | | 0.0294 | 0.903 | 0.265 | 0.151 |
| Bryophyta | Chloroplast | 21 | –0.229 | 0.158 | –0.229 | 0.158 |
| | Mitochondrial | | –0.238 | 0.140 | –0.152 | 0.354 |
| Acrogymnospermae | Chloroplast | 5 | –0.4 | 0.483 | –0.2 | 0.817 |
| | Mitochondrial | | 0.2 | 0.817 | 0.6 | 0.233 |
| Angiospermae | Chloroplast | 28 | –0.0265 | 0.860 | 0.196 | 0.150 |
| | Mitochondrial | | –0.185 | 0.174 | 0.0423 | 0.769 |
| *Loranthus* | Chloroplast | 3 | –0.333 | 1.0 | –0.333 | 1.0 |
| | Mitochondrial | | –0.333 | 1.0 | –0.333 | 1.0 |
| Chlorophyta | Chloroplast | 5 | –0.6 | 0.233 | –0.4 | 0.483 |
| | Mitochondrial | | 0.6 | 0.233 | 0.6 | 0.233 |

**Table S11. Details of independent sister-pair contrast analyses for 1st + 2nd codon sites ('nonsynonymous' sites) in nuclear and chloroplast genes, including assumption tests.** Third column reports the number of independent contrasts used in analyses after removing inappropriate contrasts. Fourth to seventh columns give Kendall's tau ($\tau$) and *p*-values for both assumption tests. Any trend in the assumption tests is indicated by bold font ($p < 0.05$).

| Taxon | Data Type | Number of Contrasts | Assumption Test 1 | | Assumption Test 2 | |
|---|---|---|---|---|---|---|
| | | | Kendall's $\tau$ | *p*-value | Kendall's $\tau$ | *p*-value |
| Monilophyta | Chloroplast | 13 | –0.0513 | 0.858 | 0.282 | 0.204 |
| | Nuclear | | 0.0769 | 0.765 | 0.410 | 0.0573 |
| Bryophyta | Chloroplast | 36 | –0.0984 | 0.409 | 0.0222 | 0.861 |
| | Nuclear | | –0.156 | 0.188 | 0.0127 | 0.925 |
| Angiospermae | Chloroplast | 7 | –0.524 | 0.136 | 0.333 | 0.381 |
| | Nuclear | | –0.143 | 0.773 | 0.0476 | 1.0 |
| Orobanchaceae | Chloroplast | 8 | –0.0714 | 0.905 | 0.214 | 0.548 |
| | Nuclear | | 0.0714 | 0.905 | 0.143 | 0.720 |

**Table S12. Details of independent sister-pair contrasts for 3rd codon sites ('synonymous' sites) in nuclear and chloroplast genes, including assumption tests.** Third column reports the number of independent contrasts used in analyses after removing inappropriate contrasts. Fourth to seventh columns give Kendall's tau ($\tau$) and *p*-values for both assumption tests. Any trend in the assumption tests is indicated by bold font ($p < 0.05$).

| Taxon | Data Type | Number of Contrasts | Assumption Test 1 | | Assumption Test 2 | |
|---|---|---|---|---|---|---|
| | | | Kendall's $\tau$ | *p*-value | Kendall's $\tau$ | *p*-value |
| Monilophyta | Chloroplast | 11 | –0.309 | 0.218 | –0.0545 | 0.879 |
| | Nuclear | | –0.273 | 0.283 | 0.127 | 0.648 |
| Bryophyta | Chloroplast | 35 | –0.150 | 0.213 | –0.0252 | 0.844 |
| | Nuclear | | –0.133 | 0.270 | 0.0252 | 0.844 |
| Angiospermae | Chloroplast | 7 | –0.143 | 0.773 | 0.333 | 0.381 |
| | Nuclear | | –0.619 | 0.0691 | 0.0476 | 1.0 |
| Orobanchaceae | Chloroplast | 7 | –0.143 | 0.773 | –0.143 | 0.773 |
| | Nuclear | | –0.333 | 0.381 | –0.238 | 0.562 |

**Table S13. Details of independent sister-pair contrasts for 1st + 2nd codon sites ('nonsynonymous' sites) in nuclear and mitochondrial genes, including assumption tests.** Third column reports the number of independent contrasts used in analyses after removing inappropriate contrasts. Fourth to seventh columns give Kendall's tau ($\tau$) and *p*-values for both assumption tests. Any trend in the assumption tests is indicated by bold font ($p < 0.05$).

| Taxon | Data Type | Number of Contrasts | Assumption Test 1 | | Assumption Test 2 | |
|---|---|---|---|---|---|---|
| | | | Kendall's $\tau$ | *p*-value | Kendall's $\tau$ | *p*-value |
| Monilophyta | Mitochondrial | 9 | –0.278 | 0.359 | –0.111 | 0.761 |
| | Nuclear | | –0.167 | 0.612 | 0.278 | 0.359 |
| Bryophyta | Mitochondrial | 19 | –0.251 | 0.143 | –0.240 | 0.164 |
| | Nuclear | | –0.193 | 0.267 | –0.0877 | 0.629 |

**Table S14. Details of independent sister-pair contrasts for 3rd codon sites ('synonymous' sites) in nuclear and mitochondrial genes, including assumption tests.** Third column reports the number of independent contrasts used in analyses after removing inappropriate contrasts. Fourth to seventh columns give Kendall's tau ($\tau$) and *p*-values for both assumption tests. Any trend in the assumption tests is indicated by bold font ($p < 0.05$).

| Taxon | Data Type | Number of Contrasts | Assumption Test 1 | | Assumption Test 2 | |
|---|---|---|---|---|---|---|
| | | | Kendall's $\tau$ | *p*-value | Kendall's $\tau$ | *p*-value |
| Monilophyta | Mitochondrial | 12 | –0.273 | 0.250 | –0.0909 | 0.737 |
| | Nuclear | | –0.394 | 0.0863 | 0 | 1.0 |
| Bryophyta | Mitochondrial | 20 | –0.0421 | 0.823 | 0 | 1.0 |
| | Nuclear | | –0.263 | 0.113 | –0.126 | 0.461 |

**Table S15. Details of independent sister-pair contrasts for 1st + 2nd codon sites ('nonsynonymous' sites) in chloroplast and mitochondrial genes, including assumption tests.** Third column reports the number of independent contrasts used in analyses after removing inappropriate contrasts. Fourth to seventh columns give Kendall's tau ($\tau$) and *p*-values for both assumption tests. Any trend in the assumption tests is indicated by bold font ($p < 0.05$).

| Taxon | Data Type | Number of Contrasts | Assumption Test 1 | | Assumption Test 2 | |
|---|---|---|---|---|---|---|
| | | | **Kendall's $\tau$** | ***p*-value** | **Kendall's $\tau$** | ***p*-value** |
| Monilophyta | Chloroplast | 17 | –0.0588 | 0.777 | 0.265 | 0.151 |
| | Mitochondrial | | 0.0147 | 0.968 | 0.0882 | 0.655 |
| Bryophyta | Chloroplast | 19 | –0.193 | 0.267 | –0.146 | 0.406 |
| | Mitochondrial | | –0.251 | 0.143 | –0.240 | 0.164 |
| Angiospermae | Chloroplast | 25 | 0.0467 | 0.764 | 0.213 | 0.142 |
| | Mitochondrial | | –0.16 | 0.276 | 0.06 | 0.694 |

**Table S16. Details of independent sister-pair contrasts for 3rd codon sites ('synonymous' sites) in chloroplast and mitochondrial genes, including assumption tests.** Third column reports the number of independent contrasts used in analyses after removing inappropriate contrasts. Fourth to seventh columns give Kendall's tau ($\tau$) and *p*-values for both assumption tests. Any trend in the assumption tests is indicated by bold font ($p < 0.05$).

| Taxon | Data Type | Number of Contrasts | Assumption Test 1 | | Assumption Test 2 | |
|---|---|---|---|---|---|---|
| | | | **Kendall's $\tau$** | ***p*-value** | **Kendall's $\tau$** | ***p*-value** |
| Monilophyta | Chloroplast | 15 | –0.276 | 0.169 | 0.0286 | 0.923 |
| | Mitochondrial | | 0.0857 | 0.697 | 0.276 | 0.169 |
| Bryophyta | Chloroplast | 15 | –0.105 | 0.627 | –0.0286 | 0.923 |
| | Mitochondrial | | 0.181 | 0.380 | 0.257 | 0.202 |
| Angiospermae | Chloroplast | 30 | –0.228 | 0.0803 | 0.126 | 0.338 |
| | Mitochondrial | | –0.177 | 0.177 | 0.0345 | 0.805 |
